# Neutrophils in the brain are sources of neuroprotective molecules and demonstrate functional heterogeneity during chronic *Toxoplasma gondii* infection

**DOI:** 10.1101/2022.08.12.503720

**Authors:** Kristina V. Bergersen, Bill Kavvathas, Clement David, Byron D. Ford, Emma H. Wilson

## Abstract

Infection with the protozoan parasite *Toxoplasma gondii* leads to the formation of lifelong cysts in neurons of the brain that can have devastating consequences in the immunocompromised. However, despite the establishment of a chronic inflammatory state and infection-induced neurological changes, there are limited signs of clinical neuropathology resulting in an asymptomatic infection in the immunocompetent. This suggests the work of neuroprotective mechanisms to prevent clinical manifestations of disease. However, such sources of neuroprotection during infection remain largely unknown. This study identifies a population of neutrophils chronically present in the brain during Toxoplasma infection that express the neuroprotective molecules NRG-1, ErbB4, and MSR1. Further phenotyping of this population via flow cytometry and singe-cell RNA sequencing reveals two distinct subsets of neutrophils based on age that display functional heterogeneity. This includes cells transcriptionally prepared to function both as anti-parasitic effector cells and in a more alternative protective manner. Chronic depletion of neutrophils results in increased parasite burden and infection-induced vascular pathology. Lack of neutrophils during chronic infection also deleteriously affects neuronal regeneration and repair mechanisms. In conclusion, this work identifies and demonstrates a functionally diverse chronic neutrophil population that plays a dynamic role in controlling infection outcome in the CNS by balancing classical responses with neuroprotective functions.

**Author Summary:** The predominantly asymptomatic nature of chronic *Toxoplasma gondii* infection despite the life-long infection of neurons suggests that there are neuroprotective mechanisms at work in the brain to maintain homeostasis and integrity. This study identifies neutrophils, normally considered a first-responding innate immune cell, as a prominent source of neuroprotective molecules during *Toxoplasma* infection. Aged neutrophils in the brain exhibit an ability to be functionally flexible expressing signatures of classical proinflammatory responses; and neuroprotective, pro-angiogenic indicators. Lack of neutrophils during chronic infection leads to increased parasite burden, increased vascular damage, and decreased neuronal regeneration. We conclude that chronic brain neutrophils are a functionally dynamic population and a source of neuroprotection during infection and suggest that this is a potentially novel target to promote brain tissue repair without compromising anti-microbial activity.

## Introduction

Ingestion of the protozoan parasite *Toxoplasma gondii* leads to a lifelong infection characterized by the formation of cysts inside neurons of the brain. Prevalence rates vary, however it is estimated that one third of the world’s population is infected with Toxoplasma [1]. While infection is often asymptomatic due to a robust host immune response, an immunocompromised state can result in parasite reactivation in the brain, the onset of Toxoplasmic encephalitis, and death [2]. Presence of the parasite in the brain leads to the recruitment of both innate and adaptive immune cells which are required to control parasite replication and maintain a balanced tissue environment [3-5].

During chronic infection, despite cyst-containing neurons and brain inflammation caused by the recruitment of peripheral immune cells, Toxoplasma causes few clinical symptoms in the immunocompetent host. However, underlying changes in neurochemistry and significant changes in the health of neurons [6, 7] suggest a constant need to balance inflammation and protect against neuropathology. Recent work in our lab to determine Toxoplasma-induced changes in immunological and neurological transcripts in the brain [8] supports these neurological changes but also reveals potential neuroprotective pathways that remain poorly understood.

The concept of neuroprotection and resolution of inflammation has been investigated in models of central nervous system (CNS) injury such as ischemic stroke and spinal cord injury and during experimental cerebral malaria (ECM) infection. In these models, neuronal integrity is maintained via signaling of the endogenous ligand Neuregulin 1 (NRG-1) through its main receptor ErbB4 [9, 10]. Resolution of neuropathology in a model of ischemic stroke is further supported by the clearance of damage signals, debris, and revascularization by macrophage scavenger receptor 1 (MSR1) [11]. Transcripts of this scavenger receptor are also upregulated during Toxoplasma infection in a step-wise manner correlating with increasing chronicity [8]. While the expression and function of these neuroprotective molecules contribute to neuroprotection in the above-mentioned diseases, their cellular sources and their functions in protecting against Toxoplasma infection and pathology remain unknown.

In recent years, research has brought recognition that the function of neutrophils is not simply to act as a first responder but can also be critical throughout immune responses including during chronic inflammation. Two recent independent research groups have identified a small population of these cells present in the brain between 2- and 4-weeks post infection (wpi) [5, 12]. Neutrophils, commonly thought of as one of the earliest host responses to infection, are one of the primary cell types responsible for the recruitment of other innate immune cells, and their presence is vital for control of acute infection [3, 13-15]. However, this discovery of neutrophils during chronic infection suggests a more versatile role for these cells. Previous literature focusing on potential alternative roles for neutrophils in non-Toxoplasma models describes two basic classes of these cells based on their differential expression of markers such as CD15 and CD49d among others [16-18]. While these earlier studies introduce the idea of alternative neutrophil functions, more recent works investigating neutrophils in homeostasis, tissue injury, and non-Toxoplasma models of infection have demonstrated a much broader range of previously undescribed heterogeneity and functions among these cells [19, 20]. Therefore, the presence of neutrophils during chronic T. gondii, may point to an important yet undefined role for these cells in the brain during chronic infection.

In this study, we have discovered the expression of neuroprotective molecules including NRG-1, ErbB4, and MSR1 among others by a persistent population of neutrophils in the brain during *T. gondii* infection. These cells have distinct phenotypic and transcriptomic profiles compared to peripheral neutrophils including CNS-specific signatures that define this population and point to a role in resolution of infection-induced neuropathology. Depletion of this population leads to an increase in vascular pathology and a decrease in neuronal regeneration during infection. The combined data reveal two populations of neutrophils that are capable of acting simultaneously as effector and neuroprotective cells. Such capacity for functional flexibility within these cells presents a paradigm shift in our understanding of neutrophils in the brain during infection and marks them as playing an important role in protecting against neuropathology caused by chronic Toxoplasma infection.

## Results

### Neutrophils in the CNS are sources of neuroprotective molecules during chronic infection

Our previous work has demonstrated the activation of potential neuroprotective and reparative pathways in the brain during chronic Toxoplasma infection [8], but the cellular sources of neuroprotection reman widely unknown. To determine sources of previously defined neuroprotective molecules in the CNS during chronic Toxoplasma infection, we performed flow cytometry on BMNCs and peripheral splenocytes at various stages of infection and examined the expression of NRG-1, ErbB4, and MSR1 (**Figure 1**).

**Figure 1.**
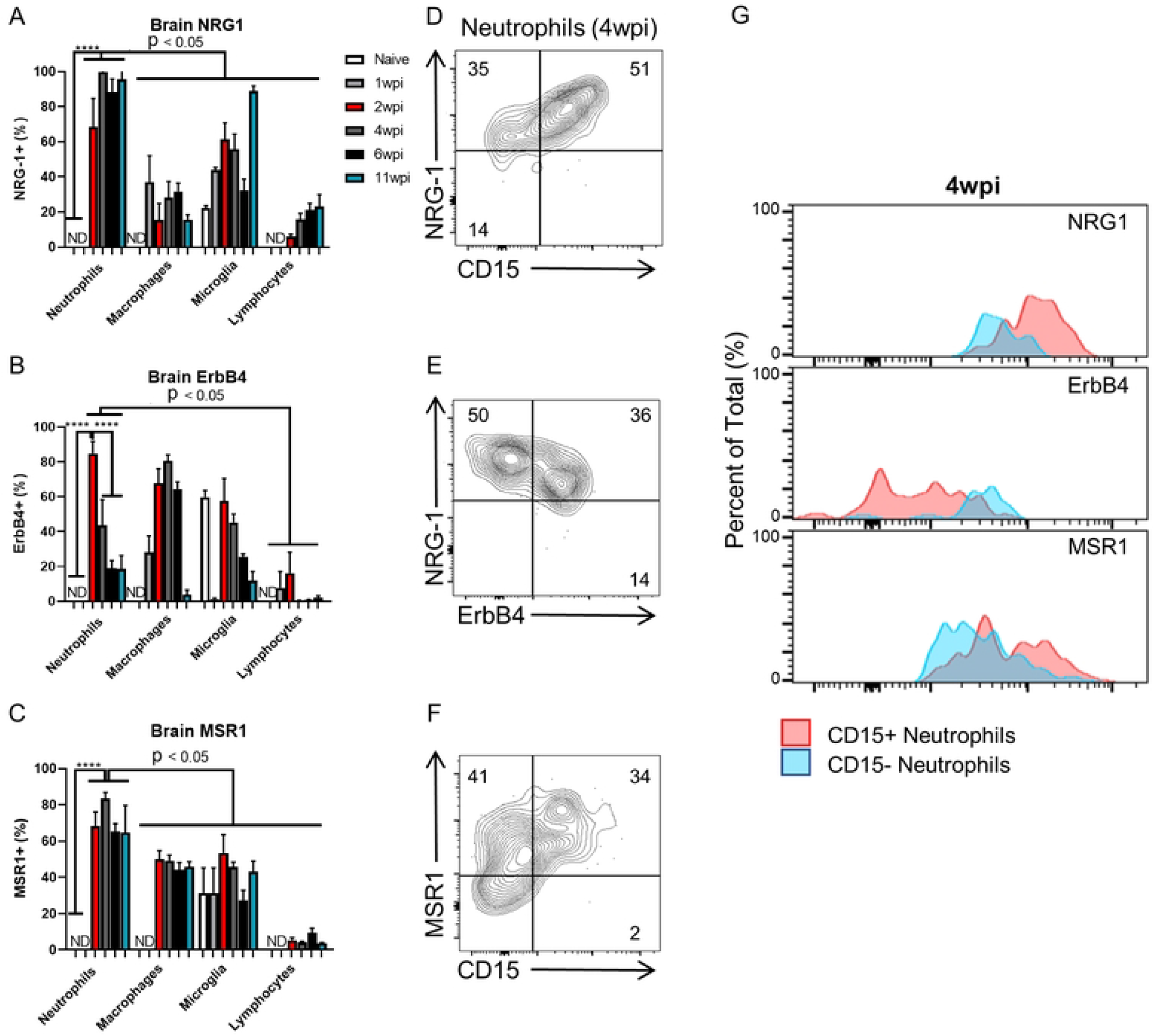
Infiltrating neutrophils in the CNS are sources of neuroprotective molecules during chronic infection. C57BL/6J mice were infected intraperitoneally with 10 *T. gondii* cysts (n=4 per time point) or injected with PBS as a control (n=3) and analyzed for immune cells in the brain and expression of neuroprotective molecules at both acute (1wpi) and chronic (2, 4, 6, and 11wpi) time points via flow cytometry. A-C) Time course quantification of expression frequencies (shown by percent of positive cells) of NRG1 (A), ErbB4 (B), and MSR1 (C) by brain mononuclear cells (BMNCs). D-F) Representative flow plots of NRG-1 (D), ErbB4 (E), and MSR1 (F) expression by neutrophils at 4wpi. Neutrophil subsets are identified based on expression of the integrin CD15 and ErbB4. Numbers on flow plots represent the average percentage of expression (n=4). G) Expression overlap of neuroprotective molecules distinguished by CD15+ (pink) and CD15- (blue) neutrophils. For all graphs, ****=P < 0.0001 (remaining significance indicated as p<0.05); significance determined via One-way ANOVA, and error bars indicate SD. Experiments were repeated 2-3 times to confirm consistency of results.

In addition to analysis of T cells, microglia and macrophages within the brain and as previously demonstrated by the Dunay lab and colleagues [5], a small but well-defined population of neutrophils was seen in the brain during chronic infection based on their expression of Ly6G and CD11b (**Supplemental Figure 1A**). Neutrophils were classified as fully differentiated and mature based on their expression of CD11b, Ly6G, MHC I, and the lack of the proliferative marker Ki67 (**Supp. Fig. 1B**). To determine the size and duration of this population over the course of infection, neutrophil percentages and absolute cell numbers were quantified from the brain at each time point. Results demonstrated consistency of this population during chronic infection after a slight decrease after 2wpi (**Supplemental Figure 2A**). Previous studies investigating the role of infiltrating inflammatory monocytes in the brain during chronic Toxoplasma infection have found infection-specific localization to the olfactory tubercle suggesting location-specific functions of these cells [12]. To determine if our neutrophil population was recruited to a particular part of the CNS, brains were perfused and the cerebellum, frontal cortex and mid-brain were dissected, anatomical areas that are easily identified and isolated. Although the percentage of neutrophils was higher in the cerebellum, neutrophils showed little localization and instead show broad disbursement throughout the brain (**Supplemental Fig. 2B)**. These results demonstrate a persistent broadly disbursed chronic brain neutrophil population.

Strikingly, flow cytometry analysis revealed that these neutrophils were consistent, significant and almost uniformly cellular sources of NRG-1 (>60% of neutrophils). A large percentage were also expressing ErbB4 (20-60% of neutrophils), and MSR1 (>60% of neutrophils) from 2wpi-11wpi (**Fig. 1A-C**). Infiltrating macrophages and resident microglia were also sources of these molecules, NRG-1 (10-30% of macrophages, 40-90% of microglia, **Fig. 1A**), ErbB4 (>60% of macrophages, 10-60% of microglia, **Fib. 1B**), and MSR1 (>40% of macrophages and microglia, **Fig. 1C**) throughout infection. Adaptive T cells did not express ErbB4 nor MSR1 at any time point, however a subpopulation (10-20%) did express NRG-1 which increased over the course of infection (**Fig. 1B**). This data demonstrates that in contrast to other immune cells, neutrophils (>60%) are consistent cellular sources of the neuroprotective molecules NRG-1, ErbB4, and MSR1 in the brain during infection.

Neutrophils were composed of two main subtypes based on their expression of the integrin CD15 (**Fig. 1D-F**). This integrin aids in migration into tissues and has previously been used to identify broad “classical” (CD15+) and “alternative” (CD15-) categories of neutrophils [18] with classical neutrophils primed as effector cells for pathogen killing and alternative neutrophils associated with healing, tissue remodeling, and vascular repair. At 4wpi, more than 80% of neutrophils were NRG-1+ and approximately 60% of these NRG-1+ cells expressed CD15, with high expression of NRG-1 correlating with expression of CD15 (**Fig. 1D**). These two neutrophil subtypes were further distinguished by their differential expression of the NRG-1 receptor, ErbB4. Alternative neutrophils that did not express CD15, were more likely to express ErbB4 and therefore be responsive to neuregulin (**Fig. 1E, G**). The expression of MSR1 was more diverse and independent of CD15 expression (**Fig. 1F**). Collectively, these results identify neutrophils as a persistent broadly disbursed cell population in the brain during chronic Toxoplasma infection. These cells are consistent cellular sources of neuroprotective molecules and are composed of 2 main subtypes based on their differential expression of CD15 and ErbB4.

### Protein analysis of chronic brain neutrophils reveals a distinct phenotypic profile

To determine if the expression of NRG-1, ErbB4, and MSR1 by brain neutrophils defined a CNS-specific protective neutrophil phenotype, cells were further analyzed for proteins known to play a role in neutrophil function. Previously identified proteins including MMP9, VEGF, CD62-L, and CXCR4 play a role in alternative/protective neutrophil functions including tissue repair and angiogenesis, but these functions have been observed predominantly in the periphery [5, 19]. To determine the expression of these molecules previously associated with protective functions by our brain neutrophils, BMNCs and peripheral splenocytes were incubated with antibodies to Ly6G, CD11b, CD15, NRG-1, ErbB4, MSR1, MMP9, VEGF, CD62-L, and CXCR4, and multi-parameter flow cytometry was conducted. Neutrophils from brain and periphery were identified using previously described techniques (**Supplemental Fig. 1**), and tSNE plots of Ly6G and CD11b expression confirmed positive neutrophil phenotype (**Supplemental Fig. 3**). When neutrophils from the brain were compared to those from the spleen, tSNE plots of all proteins analyzed revealed mostly non-overlapping populations (**Fig. 2 A**).

**Figure 2.**
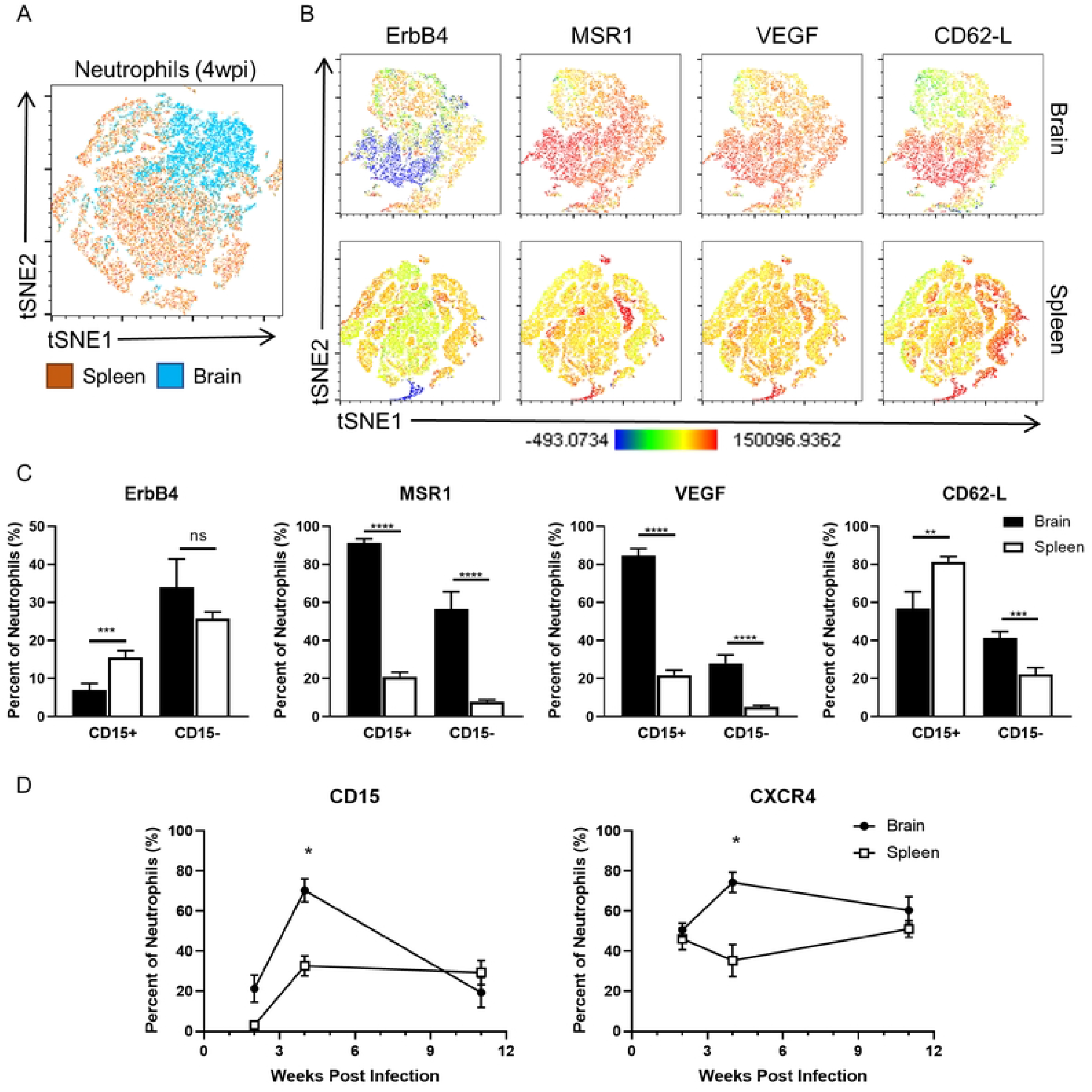
Protein analysis of chronic brain neutrophils reveals a distinct phenotypic profile. C57BL/6J mice were infected intraperitoneally with 10 *T. gondii* cysts (n=4 per time point), and neutrophil phenotypic profiles from brain and spleen were evaluated at different chronic (2, 4, and 11wpi) time points via flow cytometry. Neutrophils from brain and spleen were identified based on expression of CD11b and Ly6G (Supplemental Figure 1). A) tSNE plot of concatenated brain and spleen neutrophils at 4wpi. B) tSNE plots of selected alternative molecules from concatenated brain (top) and spleen (bottom) neutrophils at 4wpi. tSNE plot scale shows populations with low expression (blue) to high expression (red) of molecules. C) Flow cytometry quantification of selected alternative molecules by neutrophils in brain and spleen based on CD15 expression at 4wpi. ** = p< 0.01, *** = p< 0.001, **** = p<0.0001. D) Flow cytometry quantification of CD15 and CXCR4 expression by brain and spleen neutrophils at each time point during infection (2, 4, and 11wpi). * = p<0.05. For all graphs, significance determined via unpaired student t-test, and error bars indicate SD.

Expression of individual targeted proteins was evaluated to discern phenotypical differences betweenbrain and splenic neutrophils over the course of infection (**Supplemental Fig. 4**). Striking visual differences between brain and splenic neutrophils were seen in expression of ErbB4, MSR1, VEGF, and CD62-L at 4wpi (**Fig. 2B**), with neutrophils from the brain uniformly expressing higher levels of MSR1 and VEGF compared to those from the spleen. In addition, clustering analysis defined brain neutrophil subsets based on the differential expression of ErbB4 and CD62-L which was not observed in the spleen population. Additionally, brain neutrophils demonstrated higher percentages of MSR1+ and VEGF+ cells compared to splenic neutrophils. NRG-1 expression by these cells did not define further subsets but was expressed by all cells at 4- and 11wpi (**Supplemental Fig. 4A**). Contrastingly, splenic neutrophils demonstrated 2 distinct subsets based on NRG-1 expression (**Supplemental Fig. 4B**). Analysis of neutrophils from the blood served as another peripheral population and demonstrated similar results to those of the spleen compared to our brain population (**Supplemental Figure 5**). This data demonstrates a distinct phenotypic profile of brain neutrophils characterized by differential expression of targeted neuroprotective molecules when compared to periphery. As such, this population was defined as unique and will be referred to as “neut^B^ cells” for the duration of this manuscript.

It was thought that the distinctly neuroprotective phenotype observed in neut^B^ cells could be caused by the alteration of one specific subset (CD15+ or CD15-) upon entry into the brain. To test this hypothesis, the above tSNE plots were quantified based on CD15 positivity. Results confirmed neut^B^ expression patterns of ErbB4 and MSR1 based on CD15 positivity as expected (**Fig. 1**). Significant differences were observed between neut^B^ cells and splenic neutrophils independent of CD15 expression in the expression of ErbB4, MSR1, VEGF, and CD62-L (**Fig. 2C**). These results demonstrate that the phenotypic differences in neuroprotective molecule expression observed between neut^B^ cells and peripheral neutrophils encompass the neut^B^ population as a whole, and it is not one specific subset of these cells (CD15+/-) that drives the development of a CNS-specific profile.

In addition to being described as classical and alternative, neutrophils can also be distinguished by age and changes in migratory capability as reviewed by Peiseler and Kubes [19]. Thus, it was hypothesized that differences in migration capability and age would be apparent in neut^B^ cells compared to splenic neutrophils as chronic infection progressed. To address this, the expression of CD15 (indicative of the ability to migrate [18]) and CXCR4 (indicative of aged neutrophils [21]) was monitored over the course of chronic infection (**Fig. 2D**). The percent of CD15+ neut^B^ cells increased from 2wpi (20%) to 4wpi (>60%) and dropped back to baseline by 11wpi (20%), but CD15+ splenic neutrophils remained consistent (∼30% of whole population) from 4wpi on (**Fig. 2D, left**). Differences were also observed between CXCR4+ neut^B^ and peripheral cells; the percent of CXCR4+ neut^B^ cells increased by 10% from 2- to 4wpi and remained consistent through 11wpi while peripheral CXCR4 expression decreased by 20% from 2- to 4wpi and gradually returned to baseline by 11wpi (**Fig. 2D, right**). The most notable difference in expression of these molecules occurred at 4wpi. These results confirm the hypothesized differences in migration capability and age between neut^B^ and peripheral cells over time. Taken together, these results identify a phenotypically distinct neut^B^ population that differs from the spleen in its expression of alternative-associated proteins and demonstrates changes in migratory capability and age

### Aged resident and non-aged infiltrating subsets of *neut^B^* cells display functional heterogeneity

Having previously identified a unique phenotype and proteins associated with different neut^B^ subsets, we performed single cell RNA sequencing (scRNAseq) on sorted neut^B^ and peripheral cells to further assess tissue-dependent differences in the transcriptomic profile of neutrophils during chronic infection. To accomplish this, neutrophils were sorted from pooled brains and spleens of 4wk-infected mice based on positive CD11b and Ly6G expression. Resulting samples were aggregated following sequencing, and transcriptomic profiles of neut^B^ cells vs splenic cells were analyzed (**Figure 3 and Supplemental Figure 6**). To confirm positive neutrophil signatures in all datasets, *Ly6G* gene enrichment was evaluated in all sequenced samples (**Supplemental Fig. 7**). When comparing all aggregated samples in UMAP format, neut^B^ cells (red cells) clustered predominantly separately from splenic neutrophils (yellow = spleen from infected mouse, orange = spleen from naive mouse) and control BMNC cells (**Supplemental Figure 6**). When all sorted neutrophils were aggregated and analyzed separately, a nearly complete separation of neut^B^ cells (shades of red) from infected (shades of green) and naïve (shades of blue) spleen neutrophils was observed (**Fig. 3A**). This distinct clustering of neut^B^ cells at the transcriptional level supports CNS-dependent transcriptional changes and potential CNS-specific functions of these cells.

**Figure 3.**
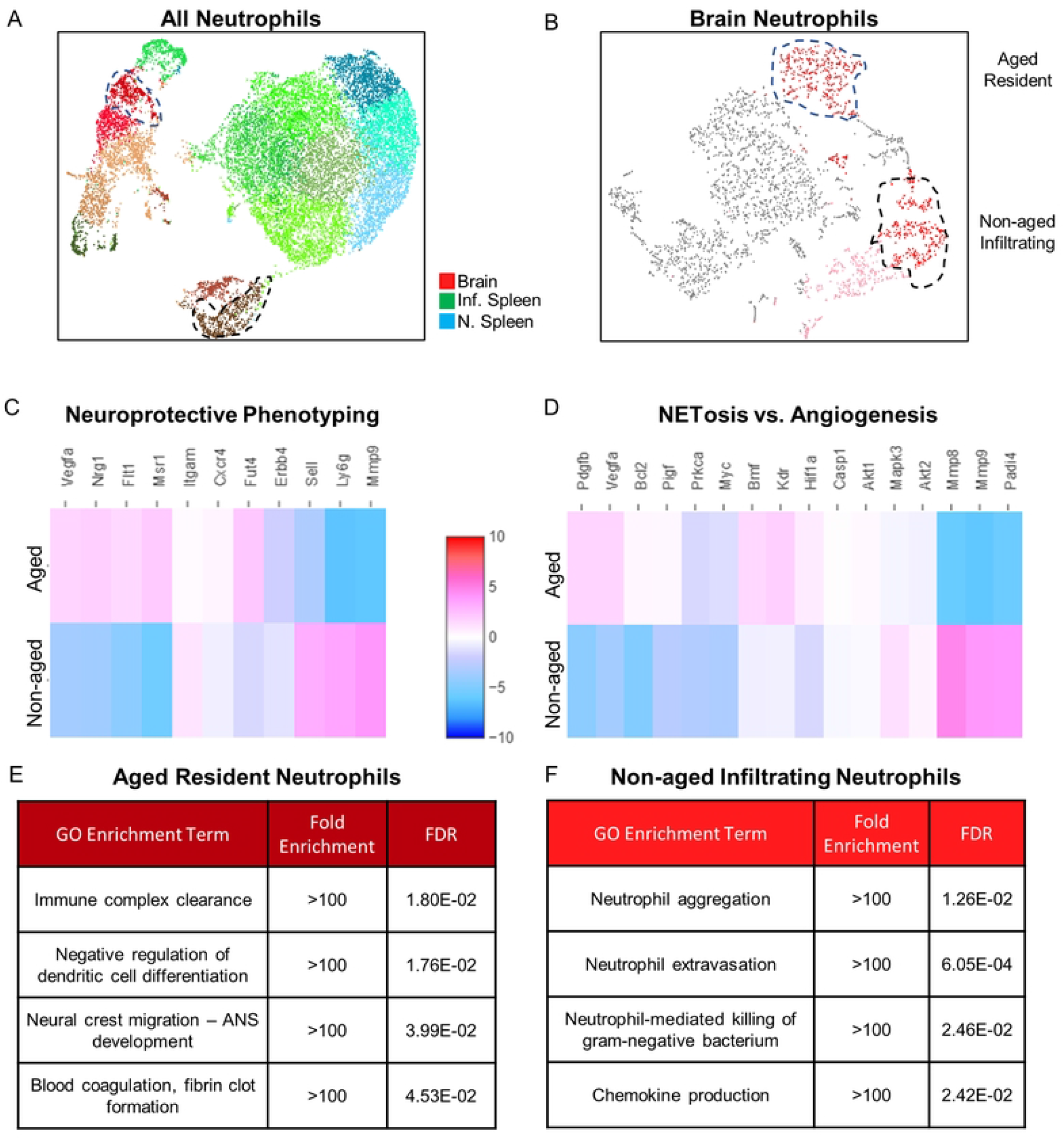
Aged resident and non-aged infiltrating subsets of neut^B^ cells display functional heterogeneity. Neutrophils from chronically infected mice were sorted from brain (n=7) and spleen (n=3) at 4wpi via flow cytometry and prepped for single cell RNA sequencing along with chronically infected BMNC (n=3) and naïve spleen (n=3) controls. Following sequencing, samples were aggregated via Loupe Browser software and also analyzed separately. A) UMAP plot of aggregated sorted neutrophil samples from Brain, Infected Spleen (Inf. Spleen), and Naïve Spleen (N. Spleen). B) UMAP plot of brain neutrophils (red = neutrophil population) identified as “Aged Resident” (blue outline, *Ly6G*+,*Fut4*+,*CXCR4*+) and “Non-aged Infiltrating” (black outline, *Ly6G*^hi^,*Fut4*^int^,*CXCR4*-) neutrophil subsets. Subsets are also identified in aggregated dataset. C-D) Heat maps of neuroprotective genes (C) and genes relating to classical (NETosis) vs alternative (angiogenesis) functions. E-F) GO enrichment terms from top 50 most significantly upregulated genes in “Aged” (E) and “Non-aged” (F) neutrophils.

scRNAseq analysis of neut^B^ cells alone resulted in 8 distinct clusters. Of these clusters, two (shades of red) were confidently identified as neutrophils based on their enrichment of *Ly6G* and *Itgam* (encodes CD11b) (**Fig. 3B**). To identify remaining cell types, targeted genes for known cell types (T cells, macrophages/microglia, inflammatory monocytes, neurons, astrocytes, and oligodendrocytes) were analyzed (**Supplemental Figure 8**). Heatmap visualization of gene expression identified several cell types: red blood cells (Cluster 5), resident CNS cells (Cluster 1 and 8), T cells (Cluster 7), and unknown/potential apoptotic cells (Cluster 2 and 3) respectively. The two identified neutrophil clusters (Clusters 4 and 6) were identified in our aggregated dataset (**Fig. 3A, blue and black outlines**) and were further classified based on enrichment of the genes *Fut4* (encodes for CD15) and *Cxcr4* based on our flow cytometry results and recent literature (reviewed by Peisler and Kubes [19]). Cluster 4 (**Fig. 3B, blue outline**) was enriched for *Ly6G, Fut4*, and *Cxcr4* and termed “Aged Resident Neutrophils.” Cluster 6 (**Fig. 3B, black outline**) was enriched for *Ly6G*, showed intermediate enrichment for *Fut4*, and was not enriched for *Cxcr4*. This cluster was named “Non-aged Infiltrating Neutrophils.” These results demonstrate two major subsets of neut^B^ cells namely “Aged” and “Non-aged” defined primarily by *Cxcr4* enrichment.

When compared directly to each other, “Neuroprotective Phenotyping” heatmap analysis demonstrated enrichment for *Vegfa, Nrg1, Flt1* (encodes for VEGF receptor), and *Msr1* in Aged-resident cells while Non-aged Infiltrating cells showed enrichment for *Sell* (encodes for CD62-L) and *Mmp9* and had increased *ErbB4* compared to the Aged Resident subset (**Fig. 3C**). Heatmap comparison analysis looking at gene enrichment corresponding to classical NETosis and alternative angiogenesis demonstrated enrichment of genes relating to both NETosis and angiogenesis in each subtype indicating functional heterogeneity of these cells (**Fig. 3D**). Aged Resident cells demonstrated enrichment for *Bmf* (NETosis gene) and the angiogenic genes *Pdgfb, Vegfa, Kdr*, and *Hif1a*, and non-aged Infiltrating cells were enriched for the NETosis-associated genes *Mapk3* and *Padi4* and the angiogenic genes *Mmp8* and *Mmp9*. Specific indicators of apoptosis were also examined including *Bcl2, Casp1, Akt1*, and *Akt2* and showed no significant enrichment in either subset. These results show differential enrichment for neuroprotective-associated genes and function-related genes between neut^B^ cells which demonstrates functional heterogeneity.

GO analysis of the top 50 most enriched genes in each neut^B^ subset showed different upregulation of GO terms (**Fig. 3E-F**). Aged Resident cells were most enriched for terms relating to negative regulation and development (**Fig. 3E**). In contrast, non-aged infiltrating cells were enriched for terms associated with classical neutrophils functions (**Fig. 3F**). Taken together, these sequencing results demonstrate a brain-specific transcriptomic profile of neut^B^ cells which can be separated into two major subsets that display neuroprotective characteristics and functional heterogeneity.

### Neutrophils during chronic infection are required to control parasite burden

To determine the requirement of these functionally flexible neut^B^ cells for control of chronic CNS infection, neutrophils were systemically depleted over the course of 2 weeks at starting at 4wpi using a neutralizing Ly6G mAb. Resulting brain and splenic neutrophil numbers, overall brain parasite and cyst burden, and brain pathology were evaluated (**Figure 4**). Neutrophils were again defined as CD11b+Ly6G (Clone 1A8)+ for gating strategy (**Supplemental Fig. 1A**). Flow cytometry analysis demonstrated >90% depletion of neutrophils in brain and spleen (**Supplemental Fig. 9A**). Absolute numbers of neutrophils in both brain and spleen were significantly lower following depletion compared to infected controls indicating successful neutrophil depletion (**Fig. 4A**). Neutrophils in the brain decreased ∼4-fold following systemic depletion, and peripheral spleen neutrophils similarly decreased ∼5-fold. To confirm that this depletion method was neutrophil-specific and did not affect other infiltrating immune cells, macrophage numbers in brain were also quantified post-depletion and were found to be unaffected (**Supplemental Fig. 9B**).

**Figure 4.**
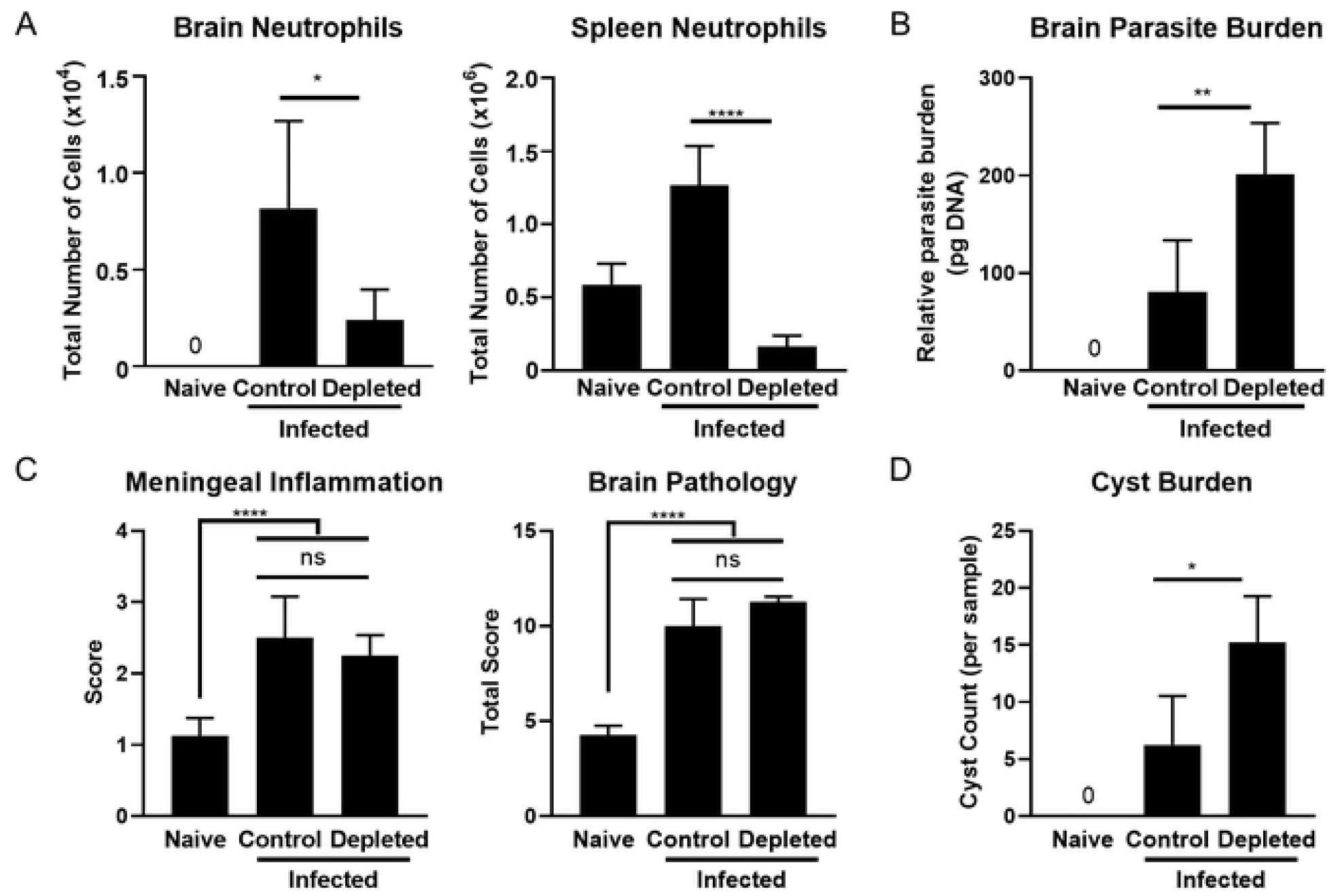
Depletion of neutrophils during chronic infection leads to increased parasite and cyst burden. C57BL/6J mice were infected intraperitoneally with 10 *T. gondii* cysts or injected with PBS as a control (n=5 per group), and a cohort of infected mice received neutralizing Ly6G mAb treatment at 4wpi for 2 weeks to deplete neutrophils. A) Total quantified numbers of neutrophils in brain (left) and spleen (right) following Ly6G mAb treatment as determined via flow cytometry. B) Quantified parasite burden from whole brain DNA via RT-PCR using *T. gondii* B1 gene (n=5/time point). C) Histopathological analysis of brains using pre-defined scoring system (see Methods). D) Cyst burden quantified via direct counting from H&E-stained slides. *= p< 0.05, **= p< 0.01, ****= p< 0.0001; significance determined via unpaired student t-test, and error bars indicate SD. Experiments were repeated 2-3 times to confirm consistency of results.

To test the role of neutrophils in parasite control and pathology prevention during chronic infection, histology and parasite burden quantification was performed on naïve, infected control, and infected neutrophil-depleted brains. Toxoplasma *B1* gene analysis demonstrated significantly elevated parasite burden (>2-fold increase) in the brain following neutrophil depletion(**Fig. 4B**). This was specific to the brain as peripheral liver parasite burden remained low but unchanged (**Supplemental Fig. 9C**). Histological analysis quantified meningeal inflammation, overall brain pathology, and cyst burden, and no significant difference was observed in inflammation or overall brain pathology (**Fig. 4C**). However, cyst burden was significantly higher in neutrophil-depleted mice compared to infected controls supporting our parasite burden results (**Fig. 4D**). Collectively, these experiments demonstrate significant increases in both parasite and cyst burden in the absence of chronic brain neutrophils.

### Lack of neut^B^ cells leads to increased blood brain barrier permeability

It is known that Toxoplasma employs various mechanisms to enter into the brain environment including transmigration through cells in the blood brain barrier (BBB) during the early stages of infection [22]. Supporting this previous finding, Evans Blue (EB) staining of infected brains during acute and chronic stages of infection demonstrated an initial increase in EB disbursement into the brain indicative of increased BBB permeability (**Fig. 5A, left**). This dissipated through 2wpi and was then present again at 4- and 6wpi. Quantification of these results confirmed significant increases in EB staining intensity at 3dpi and at the chronic time points of 4- and 6wpi (**Fig. 5A, right**). The enrichment of pro-angiogenic and vascular repair-associated genes in neut^B^ cells (**Figure 3**) led us to hypothesize that neut^B^ cells play a role in angiogenesis and vascular repair. To test this, we sought to determine the effect of neutrophil depletion on angiogenesis, BBB permeability, and vascular repair (**Fig. 5B-C**). Blinded quantification of blood vessels from naïve, infected control, and infected neutrophil-depleted brains demonstrated significantly more blood vessels counted in infected control and neutrophil-depleted brains compared to naïve controls (**Fig. 5B**) which suggests damage to followed by repair of the vasculature in the brain as demonstrated in **Fig. 5A**. However, total blood vessels quantified from neutrophil-depleted brains did not differ significantly from infected control brains suggesting that neut^B^ cells do not play a role in angiogenesis. When BBB permeability was investigated via scoring of perivascular cuffing severity following neutrophil depletion, neutrophil-depleted brains demonstrated increased perivascular cuffing severity compared to infected and naïve controls (**Fig. 5C**). These results demonstrate that while a lack of neut^B^ cells does not affect vascular angiogenesis, it does lead to increased BBB permeability.

**Figure 5.**
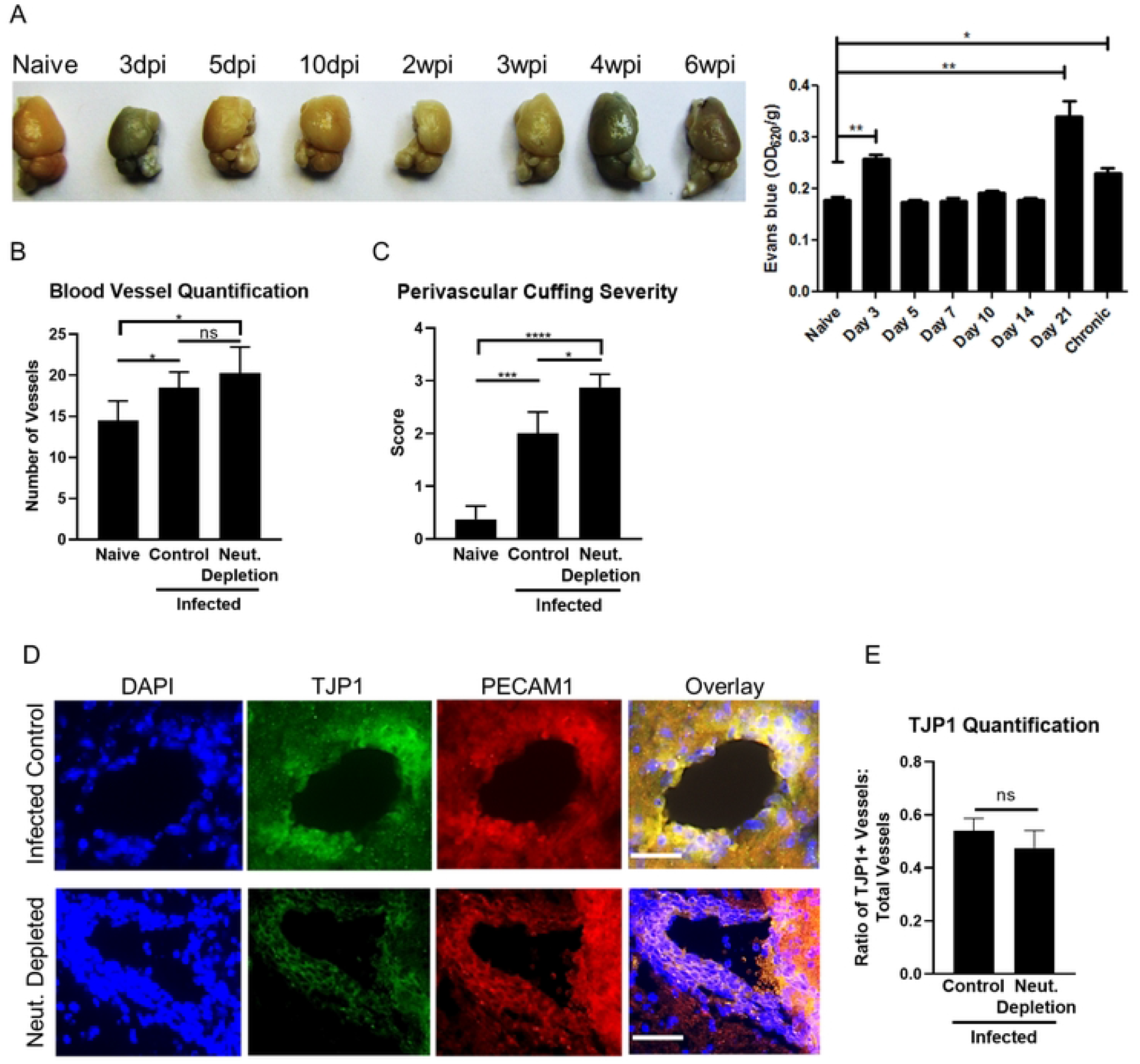
Lack of chronic brain neutrophils leads to increased vascular pathology. C57BL/6J mice were infected intraperitoneally with 10 *T. gondii* cysts or injected with PBS as a control (n=5 per group), and a cohort of infected mice received neutralizing Ly6G mAb treatment at 4wpi for 2 weeks to deplete neutrophils. A) Evans Blue staining images (left) and quantification (right) of naïve and infected mouse brain at varying stages of infection. B-C) Blinded histological analysis of brains quantifying total blood vessels counted from a minimum of 5 randomized fields of view (FOV) (B) and perivascular cuffing severity (C) using pre-defined scoring system (see Methods). D-E) Immunofluorescence images (D) and quantification (E) of infected control and neutrophil depleted brains, 40x images. Blue = DAPI, Green = TJP1, Red = PECAM1, Scale bar = 25µm. *=P < 0.05, **=P < 0.01, ****=P < 0.0001; significance determined via 1-way ANOVA and unpaired student t-test, and error bars indicate SD.

Because a lack of neut^B^ cells led to increased BBB permeability which is often detrimental to the host if not controlled, it was hypothesized that neut^B^ cells play a necessary role in vascular repair attempts and vascular remodeling. To test this, infected control and neutrophil-depleted brains were incubated in antibodies for mature blood vessels (PECAM1+) and newly formed vessels (TJP1+) [23, 24], and ratios of newly formed blood vessels: total blood vessels were compared. Qualitative differences in staining for TJP1 were observed between blood vessels from infected control and neutrophil-depleted brains (**Fig. 5D**). TJP1+ vessels from neutrophil-depleted brains demonstrated staining constricted to the outer edge of the vessel (**Fig. 5D, top**) while infected control vessels exhibited more diffuse cytoplasmic staining indicating increased regeneration (**Fig. 5D, bottom**). The overall ratio of TJP1+ vessels: total vessels did not change following neutrophil depletion (**Fig. 5E**), but differences in TJP1 expression observed in **Fig. 5D** indicates a change in vascular repair capability in the absence of neutrophils. Taken together, these results demonstrate increased BBB permeability and differences in vascular repair-related protein expression in the brain in the absence of neut^B^ cells.

### Neuronal regeneration attempts during chronic infection are inhibited in the absence of neutrophils

Neutrophils have previously been identified as neuroprotective in the CNS by encouraging neuronal regeneration via promoting upregulation of GAP43 by neurons in models of optic nerve and spinal cord injury (SCI) [25, 26]. Additionally, neutrophil expression of secretory leukocyte protease inhibitor (SLPI) and neutrophil-dependent neuronal expression of SLPI has been shown to encourage regeneration of neuronal axons in models of SCI and optic nerve injury [27, 28]. To determine whether neut^B^ cells are directly neuroprotective (actively influencing neuronal function) during Toxoplasma infection, we investigated GAP43 and SLPI expression in the brain following depletion of neutrophils (**Figure 6**).

**Figure 6.**
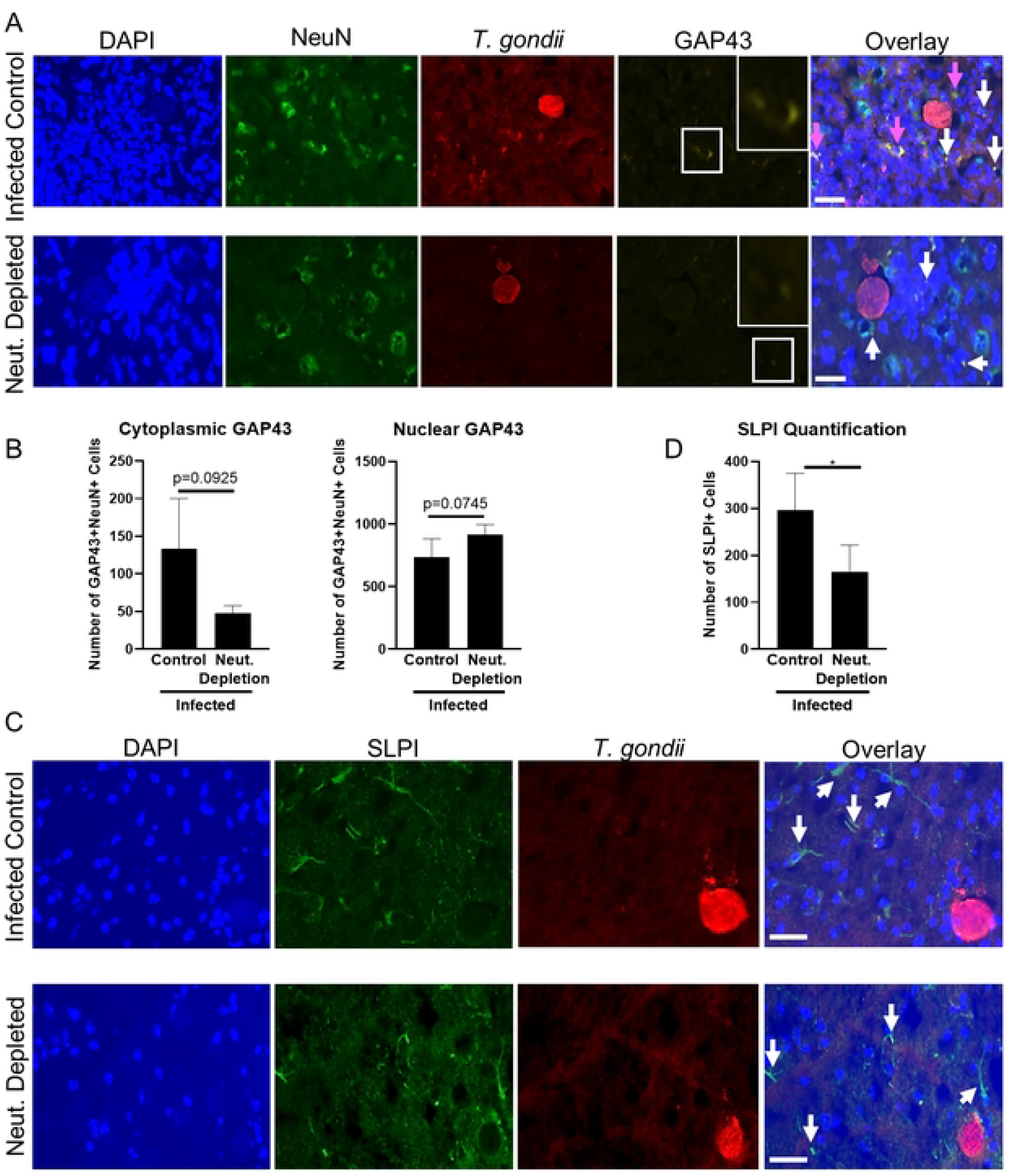
Neuronal regeneration attempts during chronic infection are inhibited in the absence of neutrophils. C57BL/6J mice were infected intraperitoneally with 10 *T. gondii* cysts or injected with PBS as a control (n=5 per group), and a cohort of infected mice received neutralizing Ly6G mAb treatment at 4wpi for 2 weeks to deplete neutrophils. A-B) Immunofluorescence images (A) and blinded quantification (B) of infected control and neutrophil depleted brains, 40x images. Blue = DAPI, Green = NeuN, Red = *T. gondii*, Yellow = GAP43, Scale bar = 25µm. White arrows indicate positive nuclear GAP43 staining, pink arrows indicate positive cytoplasmic GAP43 staining. C-D) Immunofluorescence images (C) and blinded quantification (D) of infected control and neutrophil depleted brains, 40x images. Blue = DAPI, Green = SLPI, Red = *T. gondii*, Scale bar = 25µm. White arrows indicate SLPI+ cells. *=P < 0.05, **=P < 0.01, ****=P < 0.0001; significance determined via 1-way ANOVA and unpaired student t-test, and error bars indicate SD.

Immunofluorescence imaging demonstrated visual differences in GAP43 staining between infected control and neutrophil depleted brains (**Fig. 6A**). Two different types of GAP43 expression were observed in both control and depleted tissues: 1) GAP43 staining that was restricted to the nucleus indicative of activation of GAP43 signaling but not active regeneration [1] and 2) diffuse GAP43 expression throughout the cell cytoplasm indicative of actively regenerating neuronal cells [25, 30]. Infected control brains demonstrated both obvious GAP43+ cytoplasmic and nuclear staining (**Fig. 6A, top panel, pink and white arrows**) while neutrophil-depleted brains demonstrated primarily GAP43+ nuclear staining (**Fig. 6A, bottom panel, white arrows**). When both types of GAP43 staining were blindly quantified, neutrophil-depleted brains showed a trend towards decreased cytoplasmic and increased nuclear-restricted GAP43+ staining, but these trends did not reach significance (**Fig. 6B**). These results demonstrate that neutrophils do not significantly alter neuronal regeneration during infection via the GAP43 signaling pathway.

While neutrophils were not responsible for activation of the GAP43-dependent neuronal regeneration pathway, immunofluorescence imaging of SLPI demonstrated significant differences following depletion of neutrophils (**Fig. 6C-D**). SLPI+ cell bodies and SLPI+ cell projections (shown by white arrows) were identified in both infected control and neutrophil-depleted brains (**Fig. 6C**). Based on morphology, SLPI+ cell bodies and projections were confidently identified as neurons and neuronal axons respectively and were classified as “SLPI+ cells”. When this data was blindly quantified, mice lacking neutrophils showed significantly fewer SLPI+ cells when compared to infected controls (**Fig. 6D**). These results collectively demonstrate neutrophil-dependent neuronal axon regeneration during chronic Toxoplasma infection.

## Discussion

Mechanisms of neuronal repair are critically important across a wide range of neurological and immunological diseases, yet they are poorly understood during infection where the need to control the pathogen often contradicts the traditional healing process. Toxoplasma is an infection that does not lead to severe clinical pathology despite a continuous inflammatory response in the CNS and the direct infection of neurons by the parasite. It therefore provides a useful model to address mechanisms of neuro-immune balance in the brain during chronic infection. The work here, collectively characterizes a population of chronic neutrophils in the brain during Toxoplasma infection and, in contrast to their known pro-inflammatory, anti-parasitic properties, identifies them as a source of neuroprotective molecules. These brain neutrophils are distinct from peripheral subsets and display functional heterogeneity balancing classical infection control and alternative protective mechanisms. Furthermore, we demonstrate that these neutrophils are not only required for control of parasite replication but are also required for neuronal repair during chronic infection.

Previous work has identified neutrophils and chemokine-dependent neutrophil recruitment as a vital source of protection against acute *T. gondii* infection [15, 31-33], and during the transition from acute to chronic infection in the brain [5]. Neutrophils are pre-formed sources of IL-12 [34] and that lack of IL-12 leads to a decreased IFNγ response which inhibits control of infection [35, 36]. While it is known that neutrophils are important during acute and early chronic infection, we now show that this is a highly persistent population in the brain and therefore unlikely to be a first responder population either to parasite infection nor initial invasion of the CNS. Instead, these cells express neuroprotective molecules including NRG-1, ErbB4, and MSR1 which suggest a previously undefined role of neutrophils in the modulation of neuroinflammation.

NRG-1/ErbB4 signaling-dependent neuroprotection is vital for host survival in instances of CNS damage such as ischemic stroke, spinal cord injury, epilepsy, and experimental cerebral malaria infection [9, 10, 37-39]. NRG-1 production has been broadly characterized predominantly in CNS-resident cells, specifically neurons and microglia [40-42]. However, expression of NRG-1 by migrating peripheral immune cells is less understood, and expression by neutrophils has not been previously documented. This recruitment of NRG-1 expressing immune cells may be an innate response to injury or an indication of the degree of CNS insult and therefore a need for additional peripheral support. Although NRG-1 is directly neuroprotective, the importance of MSR-1 co-expression is less clear cut. In the liver, MSR1 can activate neutrophil complement responses and worsen hepatitis. However, in the brain, it is more frequently observed in responses to DAMP signaling and can be beneficial during tissue damage [11, 43-45]. Indeed, MSR-1-mediated complement activation on neutrophils may have a different connotation within the brain, with synaptic pruning being dependent on the complement pathway. This process is part of the normal homeostasis of the brain and is required to clear non-functioning neuronal synapses. Thus, neutrophil expression of MSR1 may have a directly beneficial role on neurons by clearing damaged synapses. Therefore, the combined neutrophil expression of MSR-1 with the known neuroprotective NRG-1 and ErbB4 during chronic infection seems to support a beneficial role for MSR-1 in this context.

Neutrophils were almost uniformly positive for NRG-1 suggesting it to be a primary purpose of this population. However, it is important to note that neutrophils were not the only producers of NRG-1, and macrophages and microglia were also expressers of NRG-1, ErbB4 and MSR-1. In contrast, adaptive lymphocytes were rarely sources of these molecules. There is increasing recognition of the connection between the nervous and immune systems, however this data points to a niche for adaptive T cell immunity being primarily about controlling infection and immunomodulation and not direct neuroprotection at least via these mechanisms. Meanwhile, innate neutrophil production of neuroprotective molecules may indicate a shift in the needs of the brain once adaptive lymphocytes take over the primary anti-parasitic functions. This is supported by data demonstrating neutrophils pivoting to protective functions in the CNS following spinal cord and optic nerve injury, ischemic stroke, and retinopathy [26, 27, 46-49]. During Toxoplasma infection, the constant presence of parasite and cyst reactivation within neurons means there is likely a sustained requirement for neuroprotection, and therefore, the need for the long-term presence of a small, consistent reparative neutrophil population.

Our work not only identifies neutrophils as previously unknown sources of neuroprotective molecules but also demonstrates distinct neutrophil subsets that are specific to the brain environment. Previous work has identified two broad categories of neutrophils that differ in their functions. The first is a “classical” pro-inflammatory subset that is associated with classical macrophage activation, increased granularity, NETosis, and the production of Th1 cytokines IL-12 and TNFα [17, 18]. The second class is considered “alternative” based on anti-inflammatory cytokine expression like IL-10 and IL-4, association with alternative macrophage activation, decreased granularity, and angiogenesis and vascular repair [16, 17]. Here, we demonstrate the presence of two major neutrophil subsets in the infected brain and that this composition is different from that in the periphery suggesting that tissue-specific signals may drive the differentiation or retention of neutrophil subsets at the site of infection. Neutrophils in the brain can be differentiated into two main populations, CXCR4^hi^ aged neutrophils and non-aged CXCR4^lo^ cells [19]. Although both populations exhibit signs of functional heterogeneity, the latter non-aged population has an enhanced effector signature indicated by increased expression of genes and GO terms associated with canonical pro-inflammatory and anti-microbial functions. This supports the concept that as neutrophils enter the brain during infection, they are primarily there as an anti-microbial responsive unit. However, as they age within the brain and receive tissue-specific signals and cues, they adapt and become an enhanced neuroprotective angiogenic population. The communication between the brain and periphery that continues to draw in non-aged neutrophils even after 11wpi will be important to understand and may require parasite recrudescence and continued CCL2-dependent recruitment. In addition, a small cluster of aged neutrophils is observed in the spleen. This would either indicate differentiation of these cells in the periphery prior to migration to tissues or trafficking of tissue-educated neutrophils back into circulation and secondary lymphoid compartments. Such a population could act as an indication of continued CNS inflammation and thereby be a stimulus for new neutrophil recruitment. Whichever way these neutrophils differentiate, the question still remains as to what signals drive the brain tissue specificity of these cells and what factors encourage the differentiation into a protective phenotype. It is possible that neuronal antigens are somehow travelling to peripheral sites to initiate this process, but the mechanisms behind this subset differentiation remain to be elucidated.

Anytime there is infection or inflammation in the brain as well as in the periphery, there is likely to be a requirement for both inflammatory and repair mechanisms. Neutrophils are well-known during acute Toxoplasma infection for their production of IL-12 and their ability to kill the parasite via NETosis [33, 35] which indicate “classical” functions. In other systems, neutrophils have been defined as “alternative” and have functions that aid in angiogenesis and tissue repair as reviewed by Peisler and Kubes [19]. The presence of different neutrophil subsets in the brain during Toxoplasma infection addresses the newly acquired appreciation for neutrophil heterogeneity which can be dependent on a multitude of factors including age and tissue environment [20]. Our work suggests that despite the common urge to classify immune cells as “N1” or “N2” in terms of function, these neutrophils are capable of acting as both anti-parasitic and tissue reparative in the brain microenvironment. Thus, aged neutrophils have upregulated transcription of the effector genes *Bmf* and *Pdgfb* and also *Vegfa*, and *Kdr*, noted for their roles in tissue repair. The overlap of these seemingly contradictory functions is steadily increasing with the recognition that the production of NETs and peroxidases are also required for tissue repair as well as effector mechanisms against pathogens [27, 46, 50]. This underpins the concept of innate immune cells being as critical to tissue homeostasis as they are in initiating immune responses and may pinpoint neutrophil flexibility as important not only in Toxoplasma infection but also in all models of CNS disease.

The evidence for heterogenous functions of neutrophils in disease stems partially from their discovered roles in neuronal and angiogenic repair in both spinal cord and optic nerve injury demonstrating direct and indirect protection [26, 27, 47, 50, 51]. These functions discovered in non-infection instances may be applicable to the demonstrated evidence of damaged neurons observed during chronic Toxoplasma infection [6, 7, 52]. Additionally, the continuous latent reactivation of parasitic cysts in the brain may not only require the continuous presence of these protective neutrophils but may also be a stimulus for them to remain in the brain and differentiate into reparative subsets. As such, their consistent presence throughout chronic infection may ultimately be a signal of ongoing neuronal damage which is supported by worsening neuronal function and changes in behavior over time [6, 53] and suggests increased plasticity of neutrophils in the brain and the dependence of neutrophil function on brain microenvironment. The infection-induced changes in neuronal circuitry indicates an increased need for neuronal repair attempts as well as increased infection control. The recruitment of innate immune cells such as neutrophils may be necessary for this process as specific neuronal circuitry is required to mobilize neutrophil migration from the bone marrow [54]. Previous work has demonstrated the recruitment of innate inflammatory monocytes to the olfactory bulb of the brain perhaps to remedy olfactory-dependent behavioral changes [12]. Although there is no evidence for neutrophil localization to this area, other changes in neuronal function may be dependent on neutrophil repair of previously damaged and re-wired circuits.

Collectively, this study demonstrates neuroprotective molecule expression by a persistent neutrophil population in the brain during chronic Toxoplasma infection. The expression of these neuroprotective molecules reveals a distinct brain-specific phenotypic and transcriptomic profile of these cells which also display functional heterogeneity. Finally, these neutrophils display both direct and indirect neuroprotective functions that aid in the repair of neurons. This study reveals that not only are neutrophils continuously present throughout infection, they are also required to repair infection- and inflammation-induced damage to neurons. In conclusion, these chronic neutrophils may represent a vital component of the balanced immune response to chronic infection in the brain that *Toxoplasma gondii* is only one example of. This introduces a potential new therapeutic target for treating CNS disease without negating anti-microbial functions.

## Materials and Methods

### Animals, Parasites, and Mouse Infections/Kinetics Studies

#### Animals

All research was conducted in accordance with the Animal Welfare Act, and all efforts were made to minimize suffering. All protocols were approved by the Institutional Animal Care and Use Committee (IACUC) of the University of California, Riverside. Female 6-8 weeks old WT C57BL/6J mice were obtained from Jackson Laboratories and were maintained in a pathogen-free environment in accordance with IACUC protocols at the University of California Riverside.

#### Parasites and infections

The ME49 strain of *T. gondii* was maintained in cyst form by continuous passaging in SW and CBA/J mice. Female 6-8 week old WT C57BL/6J mice were infected with 10 ME49 cysts per mouse in 200μl of sterile 1x PBS solution via intraperitoneal (IP) injection. Naïve controls received 200µl of sterile 1x PBS solution via intraperitoneal (IP) injection.

Mice were infected as described above and sacrificed at the following acute and chronic time points: 1week post infection (wpi) (an acute time point where little cyst formation is seen in the brain), 2wpi (an early chronic time point where cysts are actively forming and immune cells are recruited), 4wpi and 6wpi (2 mid-chronic time points where cysts have been fully formed and a chronic inflammatory state is well-established), and 11wpi (a late chronic time point where increased parasite reactivation can sometimes be seen). At each time point, half brain and whole spleen were harvested for flow cytometry, and the other half brain and one lobe of liver were harvested for immunohistochemistry or parasite burden, respectively.

### Neutrophil Depletion Experiments

After 4 weeks of infection, when chronic infection was well-established, a cohort of infected mice were injected with 500µg/mouse of Ly6G depleting mouse monoclonal antibody in vehicle (sterile 1x PBS) via 200µl intraperitoneal injection every other day for 14 days according to previously published protocols [5]. A cohort of infected non-depleted mice received 200μl/mouse of vehicle (sterile 1x PBS) intraperitoneally every other day for 14 days to control for depletion injections.

### Cell Processing

Naïve and infected female C57BL/6J mice were sacrificed and perfused intracardially with 20 mL of sterile 1x PBS. Blood, spleens and half brains were harvested, and splenocytes and brain mononuclear cells (BMNCs) were processed according to previously published protocols [6, 55]. For neutrophil localization studies, whole brain was harvested and dissected into 3 broad regions (cortex, mid-brain region, and cerebellum) using brain morphology to define regions for each mouse. BMNCs were then processed as above.

### Flow Cytometry

Processed cells were diluted with FACS buffer (4g BSA, 50mg EDTA, 1L 1xPBS) to 1.0 ×10^6^ cells/ml (or all cells from respective brain regions) and transferred to FACS tubes for staining. Cells were incubated with 1:10 FC Block (BD) for 10 minutes on ice followed by fluorophore-conjugated or primary unconjugated antibodies for surface staining for 30 minutes protected from light. Cells were washed with FACS buffer solution and incubated in fluorophore-conjugated secondary antibodies as needed for 30 minutes protected from light. Fluorophore-conjugated antibodies used during surface staining are as follows: CD45 PE (Invitrogen), CD11b APCCy7 (Invitrogen), Ly6G PerCPcy5.5 (Clone 1A8, BD), CD62-L APCCy7 (BD), CD3 FITC (BD), CD4 APC (Invitrogen), CD4 APCCy7 (Invitrogen), CD8 PECy7 (Invitrogen), CD11b PerCPCy5.5 (eBioscience), CD11b APC (eBioscience), Ly6G BV510 Clone 1A8 (eBioscience), CXCR4 PECy5.5 (ThermoFisher), and MMP9 AF647 (SantaCruz Biotech). Primary unconjugated antibodies and their corresponding fluorophore-conjugated secondary antibodies are as follows: NRG-1 primary antibody (Invitrogen) with Alexafluor 647-conjugated secondary antibody (ThermoFisher); NRG-1 primary (Santa Cruz Biotech) with Alexafluor 568-conjugated secondary (ThermoFisher); MSR1 primary (Invitrogen) with Alexafluor 488-conjugated or Q Dot 655-conjugated secondary (ThermoFisher); VEGF primary (NovusBio) with Alexafluor 680-conjugated secondary (ThermoFisher); and biotinylated CD15 primary (Invitrogen) with PECy7-conjugated Streptavidin (eBioscience). Following surface staining, cells were washed, fixed in 4% PFA, and resuspended in FACS buffer. For intracellular staining, cells were spun at 1500rpm in 0.3% Saponin for 10min to permeabilize cells and then incubated with FITC-conjugated ErbB4 (Santa Cruz Biotech) in Saponin to maintain permeabilization. After incubation, cells were washed and resuspended in FACS buffer. Samples were acquired using either a BD FACS Canto II flow cytometer or NovoCyte Quanteon machine, NovoSampler Q, and NovoExpress Software at the UC Riverside core facility. Analysis was conducted using FlowJo software. For specific panels and gating strategy used, see **Supplementary Table 1** and **Supplementary Figure 1**.

### Cell Sorting and Single Cell RNA Sequencing

#### Cell Sorting

Neutrophils were sorted from brain and spleen based on CD11b/Ly6G positivity. BMNCs and splenocytes were harvested as above at 4wpi. To obtain a large enough number of neutrophils for sequencing, 3 naïve and 3 infected mice were pooled for naïve and infected neutrophil splenocyte samples, and 7 infected mice were pooled for neutrophil BMNC sample. 3 separate infected mice were pooled for an unsorted BMNC control sample. Cells were incubated in fluorescent-conjugated antibodies for CD11b APCCy7 (Invitrogen) and Ly6G PE or PerCPCy5.5 (Clone 1A8; Invitrogen), rinsed, and resuspended at 1.0 × 10^7^ concentration in FACS buffer with 10% FBS to minimize cell death during sorting. Neutrophils were sorted from pooled samples based on gating strategy of Ly6G+CD11b+ cells using a MoFlo Astrios EQ Cell Sorter at 3-4% pressure to maximize cell survival. Sorted cells were re-counted to determine viability, and a minimum of 1.0 × 10^4^ sorted cells were used for single cell sequencing.

#### 10x Sequencing and Analysis

Sorted Ly6G+CD11b+ cells and unsorted BMNC control sample were processed for single cell RNA sequencing according to 10x sample prep protocols for Steps 1-3 (Manual: CG000204 RevD); instructions were followed exactly. Time spent before loading sorted cells onto 10x Chromium controller for Step 1 was <1hr. Processed samples were analyzed after Steps 2 and 3 via Bioanalyzer by the UCR Genomics Core facility for viability and concentration before proceeding to next steps. Upon completion of the 10x protocol, samples were sent to the UC San Diego IGM Genomics Center for sequencing (500 million reads/sample).

### Immunohistochemistry

A minimum of 4 biological replicates were used for each group of mice (naïve, infected, and infected neutrophil-depleted). Following perfusion, sagittal half brains were harvested and post-fixed in 4% PFA for at least 24 hours followed by 30% sucrose for 48 hours. Brains were frozen at -80°C in optimal cutting temperature compound (OCT) and sectioned sagittally at 10µm thickness using a cryostat and charged slides.

#### H&E Staining

Briefly, H&E staining protocol was as follows: fixation of slides in 95% ethanol followed, incubation in hematoxylin followed by eosin stain for 30 seconds each (with fixation step in between), fixation in 95% and 100% ethanol, and final fixation using Citrisolv. Slides were sealed with coverslips using Cytoseal (ThermoFisher) and dried overnight. Imaging was performed using ImageJ software.

#### Histology Scoring

Slides were blinded and pathological observations in brain tissues (n=4 per group) were scored using the following criteria (lowest possible score = 4, highest possible score = 13):

1. Meningeal Inflammation Score (Scale of 1-3): 1 = little/no inflammation (maximum of 1 layer of cell nuclei present in meninges); 2 = moderate inflammation (evidence of some meningeal thickening and 1-3 layers of cell nuclei present); 3 = severe inflammation (evidence of increased overall meningeal thickening and multiple layers of cell nuclei present).
2. Perivascular Cuffing Score (Scale of 1-3): 1 = no perivascular cuffing present (little/no cell nuclei present inside blood vessels); 2 = perivascular cuffing present (evidence of blood vessel swelling and presence of multiple cell nuclei (1-2 layers of cells) inside blood vessel); 3 = severe perivascular cuffing present (very swollen blood vessels and presence of multiple cells nuclei (2-3 layers of cells) inside blood vessel, also presence of lysed blood vessels with many cell nuclei in these areas).
3. Cyst burden Score (Scale of 1-4): 1 = 0 cysts counted; 2 = <10 cysts counted for whole sample (4 brain slices); 3 = 10 cysts counted for whole sample; 4 = >10 cysts counted for whole sample.
4. Overall Tisssue Integrity Score (Scale of 1-3) – based on combined meningeal inflammation and perivascular cuffing severity: 1; = undamaged tissue (little/no meningeal inflammation and no perivascular cuffing present); 2 = moderately damaged tissue (moderate meningeal inflammation and evidence of perivascular cuffing in tissue); 3 = severely damaged tissue (severe meningeal inflammation and perivascular cuffing present).

#### Blood Vessel and Cyst Quantification

For quantification of blood vessels and cysts from H&E-stained brain sections, quantification was blinded. Total numbers of blood vessels and cysts were quantified from each whole sample (n=4 per group). Total counts per sample were calculated by adding numbers of blood vessels and cysts counted in each brain section (4 sections/sample) to account for biological variability for each group.

#### Immunofluorescence

Slides were fixed and permeabilized in 75% acetone/25% ethanol for 10 minutes at room temperature. Slides were blocked with 5% Donkey serum for 30 minutes at room temperature and incubated with primary antibodies at room temperature for 1 hour protected from light. Primary antibodies and dilutions used are listed: Rabbit NeuN (1:200, Abcam); Goat *T. gondii* (1:300, Abcam); Chicken GAP43 (1:200, Novus Biologicals); Rabbit SLPI (1:200, Novus Biologicals); and Rabbit TJP1 (1:200, Novus Biologicals). The PE-conjugated PECAM1 antibody (1:100, eBioscience) was also used. After primary incubation, slides were rinsed three times with 1x PBS for 5 minutes each wash and incubated for 1 hour protected from light at room temperature with the following secondary antibodies (1:1000 dilution, Thermofisher): Donkey anti-Rabbit Alexafluor 488, Donkey anti-Goat Alexafluor 568, Donkey anti-Chicken Alexafluor 647, and Goat anti-Rabbit Alexafluor 488. For specific panels used, see **Supplementary Table 2**. Following secondary incubation, slides were washed 3x in 1x PBS for 5 minutes each wash. Coverslips were mounted on slides using VectaShield Hardset Mounting Medium with DAPI (Vector Labs), and slides were dried overnight in the dark at room temperature. Imaging was performed using a Leica inverted DMI6000 B microscope at 40x magnification and Las-X software.

#### GAP43, SLPI, and TJP1 Quantification

Slides were blinded, and a minimum of 10 positive cells were counted per whole sample (counted from a minimum of 5 regions of interest (ROIs) per section x 4 sections/sample = minimum of 20 ROIs per sample). GAP43 positive staining was characterized as either nuclear puncta [29] or cytoplasmic [25, 30] based on previous studies. For TJP1 quantification results, the ratio of TJP1+ vessels: total vessels counted was calculated.

### Parasite Burden and B1 Gene Analysis

Parasite burden in brain and peripheral liver was quantified as previously described [8, 55]. Briefly, DNA from naïve and 2, 4, 6, and 11-week infected half brains and liver lobes of mice (n=4/group) was extracted and purified using a High Pure PCR Template Prep Kit (Roche). DNA concentration of each sample was determined via NanoDrop, and all DNA was normalized to 12.5 ng/µl before amplification. Parasite burden was measured by amplifying the B1 gene of *T. gondii* by RT-PCR.

### Statistical Analyses

All experiments were repeated a minimum of 3 times to confirm accuracy and consistency of results, and all experiments were conducted with a minimum of n=3 to account for biological variability. Statistical significance for all experiments was determined by either 2-tailed unpaired Student’s t-test or One-Way ANOVA with multiple comparisons and a p-value < 0.05 was considered statistically significant. The type of statistical test run for each experimental result is indicated in the corresponding figure legends.

#### scRNAseq Analysis

Sequencing files were analyzed using the following softwares: FileZilla, HPCC Cluster, Cell Ranger (version 5.0), and Loupe Browser (version 5.0). FastQC and MultiQC were used to evaluate the quality of sequencing reads. Sequencing reads were aligned (fastq) to the mm10 mouse reference genome for analysis, and the expression of transcripts was quantified in each cell. Low quality cells were excluded from analysis using the Normalize function during analysis. Sequencing saturation was confirmed for each sample. After quality control checks, the following cells and average gene reads per cell were obtained: Brain Neutrophils = 3,242 cells, 736 reads/cell; Infected Spleen Neutrophils = 6,804 cells, 782 reads/cell; Naïve Spleen Neutrophils = 4,813 cells, 702 reads/cell; BMNCs = 2,280 cells, 1,014 reads/cell. For differentially expressed genes (DEGs), genes with p<0.05 after comparison to control values were considered significantly enriched, and all genes discussed in this study were identified as significantly altered. Selected genes for heatmap analysis were confirmed as significantly altered (p<0.05). The top 50 most significantly enriched genes in each identified neutrophil subset were used for Gene Ontology (GO) analyses, and the “Biological Processes” option was selected for the *Mus musculus* host.

## Data Availability Statement

The data presented in this study are deposited in the Gene Expression Omnibus (GEO) repository (link: https://www.ncbi.nlm.nih.gov/geo/) and are accessible through GEO Series accession number GSE210883 (https://www.ncbi.nlm.nih.gov/geo/query/acc.cgi?acc=GSE210883).

## Acknowledgments

The authors would like to thank the members of the University of California Riverside’s Center for Glial Neuronal Interactions (CGNI) and the Genomics and School of Medicine Research COREs for all advice, use of equipment, and processing of samples where appropriate. We are also thankful to the UCSD IGM Genomics Center for sequencing of RNA samples. Lastly, we sincerely appreciate the work conducted by UCR’s IACUC and the animal husbandry provided by Leslie Karpinski, Jennifer Johnson, and Linda McCloud.

## Funding

This work was supported in part by R01DA048815; R01AI158417; and R01AI124682 to EHW, R01NS125775 and R25GM119975 to BDF, and by UCR Graduate Division to EHW.

## Authors’ contributions

KVB, CD, and EHW designed and conducted experiments; BDF provided guidance on project design; KVB, BK, and EHW analyzed data; KVB and EHW wrote the manuscript.

---

**Supplemental Figure 1. Neutrophil identification and gating strategy**. A) Representative gating strategy and gating of CD11b+Ly6G+ cells from the brain of naïve controls. Neutrophils were defined as CD11b+Ly6G+ after gating on live cells. Numerical values represent average percentages. B) Representative gating of CD11b+Ly6G+ cells from the brain of T. gondii-infected mice at the early chronic stage of infection (2wpi). Non-proliferating neutrophils were then defined as Ki67-MHC I+. This gating strategy was also used for identification of neutrophils from peripheral spleen and blood.

**Supplemental Figure 2. Protective neutrophils that persist throughout chronic infection are broadly disbursed in the brain**. C57BL/6J mice were infected intraperitoneally with 10 *T. gondii* cysts (n=4 per time point) or injected with PBS as a control (n=3) and analyzed for immune cells in the brain at both acute (1wpi) and chronic (2, 4, 6, and 11wpi) time points via flow cytometry. For location studies, mice (n=5) were sacrificed at 4wpi, brains were dissected into 3 distinct regions, and BMNCs were analyzed via flow cytometry to determine neutrophil location in the brain. A) frequencies (left) and numbers (right) of CD11b+Ly6G+ neutrophils in brain at each time point. B) Quantification of neutrophil percentages (left) and numbers (right) in each defined brain region. ** = P < 0.01, * = P < 0.05, “p-val<0.05” indicates varying degrees of significance between indicated time points; significance determined via 1-way ANOVA, and error bars indicate SD. Experiments were repeated 2-3 times to confirm consistency of results.

**Supplemental Figure 3. Confirmation of neutrophil phenotype for tSNE analysis**. C57BL/6J mice were infected intraperitoneally with 10 *T. gondii* cysts (n=4 per time point) or injected with PBS as a control (n=3), and neutrophil phenotype was confirmed via expression of Ly6G and CD11b in brain, spleen, and blood at different chronic (2, 4, and 11wpi) time points via flow cytometry. Representative tSNE plots of concatenated brain (A), spleen (B), and blood (C) neutrophils are shown. tSNE plots scale shows populations with low expression (blue) to high expression (red) of molecules.

**Supplemental Figure 4. Chronic brain neutrophils demonstrate a distinct phenoptypic profile and express a range of alternative proteins**. C57BL/6J mice were infected intraperitoneally with 10 *T. gondii* cysts (n=4 per time point) or injected with PBS as a control (n=3), and neutrophil phenotypic profiles from brain, spleen, and blood were evaluated at different chronic (2, 4, and 11wpi) time points via flow cytometry. Neutrophils from brain, spleen, and blood were identified based on expression of CD11b and Ly6G (Supplemental Figure 1). tSNE plots of concatenated brain (A-C), spleen (D-F), and blood (G-I) neutrophils are shown. tSNE plots scale shows populations with low expression (blue) to high expression (red) of molecules.

**Supplemental Figure 5. Blood neutrophil phenotype** C57BL/6J mice were infected intraperitoneally with 10 *T. gondii* cysts (n=4 per time point) or injected with PBS as a control (n=3), and neutrophil phenotypic profiles from blood were evaluated at different chronic (2, 4, and 11wpi) time points via flow cytometry. Neutrophils from blood were identified based on expression of CD11b and Ly6G. tSNE plots of concatenated blood samples at 2 (A), 4 (B), and 11wpi (C) neutrophils are shown. tSNE plots scale shows populations with low expression (blue) to high expression (red) of molecules.

**Supplemental Figure 6. ScRNAseq of peripheral and brain neutrophils during chronic infection demonstrates distinct clustering of chronic brain neutrophils**. Neutrophils from chronically infected mice were sorted from brain (n=7) and spleen (n=3) at 4wpi via flow cytometry and prepped for single cell RNA sequencing along with chronically infected BMNC (n=3) and naïve spleen (n=3) controls. Following sequencing, samples were analyzed separately and aggregated via Loupe Browser software. A) UMAP plot of all aggregated samples (BMNCs (purple), Brain Neutrophils (red), Infected Spleen Neutrophils (yellow), and Naïve Spleen Neutrophils (orange)).

**Supplemental Figure 7. Identification of neutrophil populations from scRNAseq data after sorting BMNCs based on Ly6G+CD11b+ expression**. A) Ly6G log_2_ gene expression of respective aggregated UMAPs from scRNAseq to confirm positive neutrophil phenotype. B) UMAP of Ly6G log_2_ gene expression of sorted brain neutrophils only.

**Supplemental Figure 8. Identification of cell types from sorted “Brain Neutrophil” sample**. Heat map of canonical markers of various cell types found in the brain during chronic Toxoplasma infection identified via scRNAseq analysis.

**Supplemental Figure 9. Successful and specific depletion of neutrophils after treatment with neutralizing Ly6G antibody**. A) Representative gating of CD11b+Ly6G+ neutrophils from the brain and spleen of naïve controls, infected controls, and infected neutrophil-depleted mice. Neutrophils were defined as CD11b+Ly6G+ after gating on live cells. Numerical values represent average percentages. B) Number of macrophages in brain following Ly6G depletion treatment demonstrates neutrophil-specific depletion.

**Supplementary Table 1.**
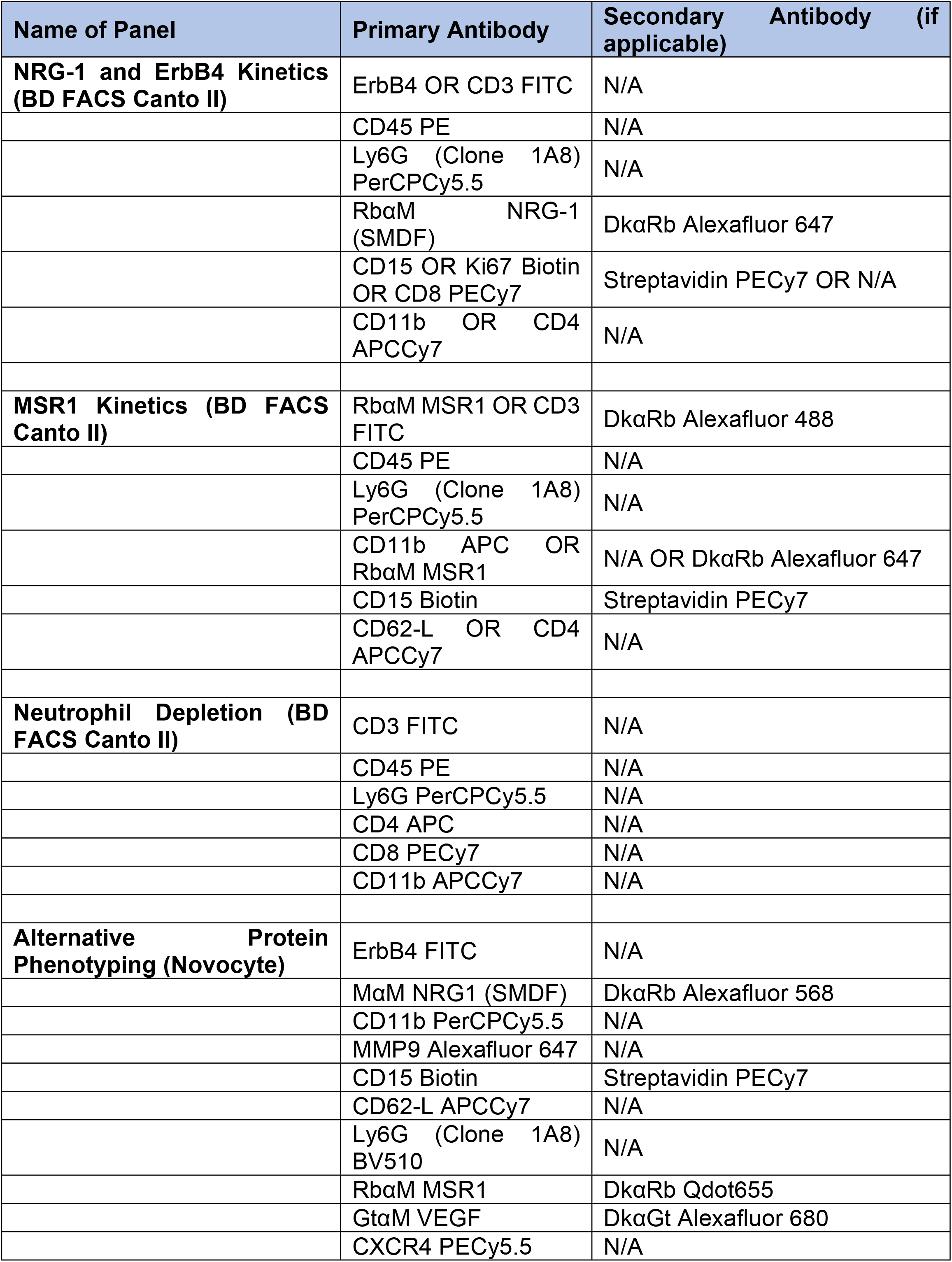
Flow cytometry antibody panels.

**Supplementary Table 2.** Immunofluorescence antibody panels.

| Name of Panel | Primary Antibody | Secondary Antibody (if applicable) |
| --- | --- | --- |
| Brain Vasculature Panel | RBαM TJP1 | GtαRb Alexafluor 488 |
|  | PECAM1 PE | N/A |
| Brain GAP43 Panel | RbαM NeuN | DkαRb Alexafluor 488 |
|  | GtαToxoplasma | DkαGt Alexafluor 568 |
|  | ChkαM GAP43 | DkαChk Alexafluor 647 |
| Brain SLPI Panel | RbαM SLPI | DkαRb Alexafluor 488 |
|  | GtαToxoplasma | DkαGt Alexafluor 568 |

## Notes

### Competing Interest Statement

I have read the journal's policy and the authors of this manuscript have the following competing interests: BF holds patents related to the work being reported without direct corporate involvement at the time.

## References

1. Montoya JG, Liesenfeld O. Toxoplasmosis. The Lancet. 2004;363(9425):1965–76. doi: https://doi.org/10.1016/S0140-6736(04)16412-X.

2. Wang Z-D, Liu H-H, Ma Z-X, Ma H-Y, Li Z-Y, Yang Z-B, et al. Toxoplasma gondii Infection in Immunocompromised Patients: A Systematic Review and Meta-Analysis. Frontiers in Microbiology. 2017;8:389.

3. McGovern KE, Wilson EH. Role of Chemokines and Trafficking of Immune Cells in Parasitic Infections. Current immunology reviews. 2013;9(3):157–68. doi: 10.2174/1573395509666131217000000. PubMed PMID: 25383073.

4. Sasai M, Yamamoto M. Innate, adaptive, and cell-autonomous immunity against Toxoplasma gondii infection. Exp Mol Med. 2019;51(12):1-10. Epub 2019/12/13. doi: 10.1038/s12276-019-0353-9. PubMed PMID: 31827072; PubMed Central PMCID: PMCPMC6906438.

5. Biswas A, French T, Düsedau HP, Mueller N, Riek-Burchardt M, Dudeck A, et al. Behavior of Neutrophil Granulocytes during Toxoplasma gondii Infection in the Central Nervous System. Frontiers in Cellular and Infection Microbiology. 2017;7:259.

6. David CN, Frias ES, Szu JI, Vieira PA, Hubbard JA, Lovelace J, et al. GLT-1-Dependent Disruption of CNS Glutamate Homeostasis and Neuronal Function by the Protozoan Parasite Toxoplasma gondii. PLOS Pathogens. 2016;12(6):e1005643. doi: 10.1371/journal.ppat.1005643.

7. Mendez OA, Flores Machado E, Lu J, Koshy AA. Injection with Toxoplasma gondii protein affects neuron health and survival. Elife. 2021;10. Epub 20210609. doi: 10.7554/eLife.67681. PubMed PMID: 34106047; PubMed Central PMCID: PMCPMC8270641.

8. Bergersen KV, Barnes A, Worth D, David C, Wilson EH. Targeted Transcriptomic Analysis of C57BL/6 and BALB/c Mice During Progressive Chronic Toxoplasma gondii Infection Reveals Changes in Host and Parasite Gene Expression Relating to Neuropathology and Resolution. Front Cell Infect Microbiol. 2021;11:645778. Epub 20210318. doi: 10.3389/fcimb.2021.645778. PubMed PMID: 33816350; PubMed Central PMCID: PMCPMC8012756.

9. Simmons LJ, Surles-Zeigler MC, Li Y, Ford GD, Newman GD, Ford BD. Regulation of inflammatory responses by neuregulin-1 in brain ischemia and microglial cells in vitro involves the NF-kappa B pathway. Journal of Neuroinflammation. 2016;13(1):237. doi: 10.1186/s12974-016-0703-7.

10. Liu M, Solomon W, Cespedes JC, Wilson NO, Ford B, Stiles JK. Neuregulin-1 attenuates experimental cerebral malaria (ECM) pathogenesis by regulating ErbB4/AKT/STAT3 signaling. Journal of Neuroinflammation. 2018;15(1):104. doi: 10.1186/s12974-018-1147-z.

11. Shichita T, Ito M, Morita R, Komai K, Noguchi Y, Ooboshi H, et al. MAFB prevents excess inflammation after ischemic stroke by accelerating clearance of damage signals through MSR1. Nature Medicine. 2017;23(6):723–32. doi: 10.1038/nm.4312.

12. Schneider CA, Figueroa Velez DX, Azevedo R, Hoover EM, Tran CJ, Lo C, et al. Imaging the dynamic recruitment of monocytes to the blood-brain barrier and specific brain regions during Toxoplasma gondii infection. Proc Natl Acad Sci U S A. 2019;116(49):24796-807. Epub 2019/11/16. doi: 10.1073/pnas.1915778116. PubMed PMID: 31727842; PubMed Central PMCID: PMCPMC6900744.

13. Debierre-Grockiego F, Moiré N, Torres Arias M, Dimier-Poisson I. Recent Advances in the Roles of Neutrophils in Toxoplasmosis. Trends Parasitol. 2020;36(12):956-8. Epub 2020/09/22. doi: 10.1016/j.pt.2020.08.007. PubMed PMID: 32952059.

14. Petri WA, Reichmann G, Walker W, Villegas Eric N, Craig L, Cai G, et al. The CD40/CD40 Ligand Interaction Is Required for Resistance to Toxoplasmic Encephalitis. Infection and Immunity. 2000;68(3):1312–8. doi: 10.1128/IAI.68.3.1312-1318.2000.

15. Bliss SK, Gavrilescu LC, Alcaraz A, Denkers EY. Neutrophil depletion during Toxoplasma gondii infection leads to impaired immunity and lethal systemic pathology. Infection and immunity. 2001;69(8):4898–905. doi: 10.1128/IAI.69.8.4898-4905.2001. PubMed PMID: 11447166.

16. Tsuda Y, Takahashi H, Kobayashi M, Hanafusa T, Herndon DN, Suzuki F. Three Different Neutrophil Subsets Exhibited in Mice with Different Susceptibilities to Infection by Methicillin-Resistant Staphylococcus aureus. Immunity. 2004;21(2):215–26. doi: https://doi.org/10.1016/j.immuni.2004.07.006.

17. Beyrau M, Bodkin J, Nourshargh S. Neutrophil heterogeneity in health and disease: A revitalized avenue in inflammation and immunity 2012. 120134 p.

18. Nakayama F, Nishihara S, Iwasaki H, Kudo T, Okubo R, Kaneko M, et al. CD15 expression in mature granulocytes is determined by alpha 1,3-fucosyltransferase IX, but in promyelocytes and monocytes by alpha 1,3-fucosyltransferase IV. J Biol Chem. 2001;276(19):16100-6. Epub 2001/03/30. doi: 10.1074/jbc.M007272200. PubMed PMID: 11278338.

19. Peiseler M, Kubes P. More friend than foe: the emerging role of neutrophils in tissue repair. The Journal of clinical investigation. 2019;129(7):2629–39. doi: 10.1172/JCI124616. PubMed PMID: 31205028.

20. Xie X, Shi Q, Wu P, Zhang X, Kambara H, Su J, et al. Single-cell transcriptome profiling reveals neutrophil heterogeneity in homeostasis and infection. Nature immunology. 2020;21(9):1119-33. Epub 2020/07/27. doi: 10.1038/s41590-020-0736-z. PubMed PMID: 32719519.

21. Casanova-Acebes M, Pitaval C, Weiss LA, Nombela-Arrieta C, Chèvre R, N AG, et al. Rhythmic modulation of the hematopoietic niche through neutrophil clearance. Cell. 2013;153(5):1025–35. doi: 10.1016/j.cell.2013.04.040. PubMed PMID: 23706740; PubMed Central PMCID: PMCPMC4128329.

22. Olivera GC, Ross EC, Peuckert C, Barragan A. Blood-brain barrier-restricted translocation of Toxoplasma gondii from cortical capillaries. Elife. 2021;10. Epub 20211208. doi: 10.7554/eLife.69182. PubMed PMID: 34877929; PubMed Central PMCID: PMCPMC8700292.

23. Liu XQ, Shao XR, Liu Y, Dong ZX, Chan SH, Shi YY, et al. Tight junction protein 1 promotes vasculature remodeling via regulating USP2/TWIST1 in bladder cancer. Oncogene. 2022;41(4):502-14. Epub 20211115. doi: 10.1038/s41388-021-02112-w. PubMed PMID: 34782718.

24. Strauss RE, Mezache L, Veeraraghavan R, Gourdie RG. The Cx43 Carboxyl-Terminal Mimetic Peptide αCT1 Protects Endothelial Barrier Function in a ZO1 Binding-Competent Manner. Biomolecules. 2021;11(8). Epub 20210812. doi: 10.3390/biom11081192. PubMed PMID: 34439858; PubMed Central PMCID: PMCPMC8393261.

25. Sas AR, Carbajal KS, Jerome AD, Menon R, Yoon C, Kalinski AL, et al. A new neutrophil subset promotes CNS neuron survival and axon regeneration. Nature Immunology. 2020;21(12):1496–505. doi: 10.1038/s41590-020-00813-0.

26. Kurimoto T, Yin Y, Habboub G, Gilbert H-Y, Li Y, Nakao S, et al. Neutrophils express oncomodulin and promote optic nerve regeneration. The Journal of neuroscience : the official journal of the Society for Neuroscience. 2013;33(37):14816–24. doi: 10.1523/JNEUROSCI.5511-12.2013. PubMed PMID: 24027282.

27. Ghasemlou N, Bouhy D, Yang J, López-Vales R, Haber M, Thuraisingam T, et al. Beneficial effects of secretory leukocyte protease inhibitor after spinal cord injury. Brain : a journal of neurology. 2010;133(Pt 1):126–38. doi: 10.1093/brain/awp304. PubMed PMID: 20047904.

28. Hannila SS, Siddiq MM, Carmel JB, Hou J, Chaudhry N, Bradley PM, et al. Secretory leukocyte protease inhibitor reverses inhibition by CNS myelin, promotes regeneration in the optic nerve, and suppresses expression of the transforming growth factor-β signaling protein Smad2. J Neurosci. 2013;33(12):5138–51. doi: 10.1523/jneurosci.5321-12.2013. PubMed PMID: 23516280; PubMed Central PMCID: PMCPMC3684282.

29. Gorup D, Bohaček I, Miličević T Pochet R, Mitrečić D. KrižJ, et al. Increased expression and colocalization of GAP43 and CASP3 after brain ischemic lesion in mouse. Neuroscience Letters. 2015;597:176–82. doi: https://doi.org/10.1016/j.neulet.2015.04.042.

30. Guarnieri S, Morabito C, Paolini C, Boncompagni S, Pilla R, Fanò-Illic G, et al. Growth associated protein 43 is expressed in skeletal muscle fibers and is localized in proximity of mitochondria and calcium release units. PLoS One. 2013;8(1):e53267. Epub 20130107. doi: 10.1371/journal.pone.0053267. PubMed PMID: 23308181; PubMed Central PMCID: PMCPMC3538766.

31. Del Rio L, Bennouna S, Salinas J, Denkers EY. CXCR2 Deficiency Confers Impaired Neutrophil Recruitment and Increased Susceptibility During <em>Toxoplasma gondii</em> Infection. The Journal of Immunology. 2001;167(11):6503.

32. Bliss SK, Marshall AJ, Zhang Y, Denkers EY. Human polymorphonuclear leukocytes produce IL-12, TNF-alpha, and the chemokines macrophage-inflammatory protein-1 alpha and - 1 beta in response to Toxoplasma gondii antigens. J Immunol. 1999;162(12):7369-75. PubMed PMID: 10358188.

33. Abi Abdallah DS, Lin C, Ball CJ, King MR, Duhamel GE, Denkers EY. Toxoplasma gondii triggers release of human and mouse neutrophil extracellular traps. Infect Immun. 2012;80(2):768-77. Epub 2011/11/23. doi: 10.1128/iai.05730-11. PubMed PMID: 22104111; PubMed Central PMCID: PMCPMC3264325.

34. Cassatella MA, Meda L, Gasperini S, D’Andrea A, Ma X, Trinchieri G. Interleukin-12 production by human polymorphonuclear leukocytes. European Journal of Immunology. 1995;25(1):1–5. doi: 10.1002/eji.1830250102.

35. Sukhumavasi W, Egan CE, Denkers EY. Mouse Neutrophils Require JNK2 MAPK for <em>Toxoplasma gondii</em>-Induced IL-12p40 and CCL2/MCP-1 Release. The Journal of Immunology. 2007;179(6):3570.

36. Gazzinelli RT, Wysocka M, Hayashi S, Denkers EY, Hieny S, Caspar P, et al. Parasite-induced IL-12 stimulates early IFN-gamma synthesis and resistance during acute infection with Toxoplasma gondii. J Immunol. 1994;153(6):2533-43. Epub 1994/09/15. PubMed PMID: 7915739.

37. Alizadeh A, Santhosh KT, Kataria H, Gounni AS, Karimi-Abdolrezaee S. Neuregulin-1 elicits a regulatory immune response following traumatic spinal cord injury. Journal of Neuroinflammation. 2018;15(1):53. doi: 10.1186/s12974-018-1093-9.

38. Tan G-H, Liu Y-Y, Hu X-L, Yin D-M, Mei L, Xiong Z-Q. Neuregulin 1 represses limbic epileptogenesis through ErbB4 in parvalbumin-expressing interneurons. Nature Neuroscience. 2011;15:258. doi: 10.1038/nn.3005

39. https://www.nature.com/articles/nn.3005#supplementary-information.

40. Cespedes JC, Liu M, Harbuzariu A, Nti A, Onyekaba J, Cespedes HW, et al. Neuregulin in Health and Disease. Int J Brain Disord Treat. 2018;4(1). Epub 20181110. doi: 10.23937/2469-5866/1410024. PubMed PMID: 31032468; PubMed Central PMCID: PMCPMC6483402.

41. Falls DL. Neuregulins and the neuromuscular system: 10 years of answers and questions. Journal of Neurocytology. 2003;32(5):619–47. doi: 10.1023/B:NEUR.0000020614.83883.be.

42. Falls DL. Neuregulins: functions, forms, and signaling strategies. Experimental Cell Research. 2003;284(1):14–30. doi: https://doi.org/10.1016/S0014-4827(02)00102-7.

43. Buonanno A, Fischbach GD. Neuregulin and ErbB receptor signaling pathways in the nervous system. Current Opinion in Neurobiology. 2001;11(3):287–96. doi: https://doi.org/10.1016/S0959-4388(00)00210-5.

44. Liu DL, Hong Z, Li JY, Yang YX, Chen C, Du JR. Phthalide derivative CD21 attenuates tissue plasminogen activator-induced hemorrhagic transformation in ischemic stroke by enhancing macrophage scavenger receptor 1-mediated DAMP (peroxiredoxin 1) clearance. J Neuroinflammation. 2021;18(1):143. Epub 20210624. doi: 10.1186/s12974-021-02170-7. PubMed PMID: 34162400; PubMed Central PMCID: PMCPMC8223381.

45. Zou X, Yang XJ, Gan YM, Liu DL, Chen C, Duan W, et al. Neuroprotective Effect of Phthalide Derivative CD21 against Ischemic Brain Injury:Involvement of MSR1 Mediated DAMP peroxiredoxin1 Clearance and TLR4 Signaling Inhibition. J Neuroimmune Pharmacol. 2021;16(2):306-17. Epub 20200414. doi: 10.1007/s11481-020-09911-0. PubMed PMID: 32291602.

46. Tang Y, Li H, Li J, Liu Y, Li Y, Zhou J, et al. Macrophage scavenger receptor 1 contributes to pathogenesis of fulminant hepatitis via neutrophil-mediated complement activation. Journal of hepatology. 2018;68(4):733-43. Epub 2017/11/14. doi: 10.1016/j.jhep.2017.11.010. PubMed PMID: 29154963.

47. Binet F, Cagnone G, Crespo-Garcia S, Hata M, Neault M, Dejda A, et al. Neutrophil extracellular traps target senescent vasculature for tissue remodeling in retinopathy. Science. 2020;369(6506). doi: 10.1126/science.aay5356. PubMed PMID: 32820093.

48. Gong Y, Koh DR. Neutrophils promote inflammatory angiogenesis via release of preformed VEGF in an in vivo corneal model. Cell Tissue Res. 2010;339(2):437-48. Epub 20091212. doi: 10.1007/s00441-009-0908-5. PubMed PMID: 20012648.

49. Hou Y, Yang D, Xiang R, Wang H, Wang X, Zhang H, et al. N2 neutrophils may participate in spontaneous recovery after transient cerebral ischemia by inhibiting ischemic neuron injury in rats. International Immunopharmacology. 2019;77:105970. doi: https://doi.org/10.1016/j.intimp.2019.105970.

50. Stirling DP, Liu S, Kubes P, Yong VW. Depletion of Ly6G/Gr-1 Leukocytes after Spinal Cord Injury in Mice Alters Wound Healing and Worsens Neurological Outcome. The Journal of Neuroscience. 2009;29(3):753. doi: 10.1523/JNEUROSCI.4918-08.2009.

51. Azcona JA, Tang S, Berry E, Zhang FF, Garvey R, Falck JR, et al. Neutrophil-derived Myeloperoxidase and Hypochlorous Acid Critically Contribute to 20-HETE Increases that Drive Post-Ischemic Angiogenesis. J Pharmacol Exp Ther. 2022. Epub 20220319. doi: 10.1124/jpet.121.001036. PubMed PMID: 35306474.

52. Christoffersson G, Vågesjö E, Vandooren J, Lidén M, Massena S, Reinert RB, et al. VEGF-A recruits a proangiogenic MMP-9-delivering neutrophil subset that induces angiogenesis in transplanted hypoxic tissue. Blood. 2012;120(23):4653-62. Epub 20120910. doi: 10.1182/blood-2012-04-421040. PubMed PMID: 22966168; PubMed Central PMCID: PMCPMC3512240.

53. Brooks JM, Carrillo GL, Su J, Lindsay DS, Fox MA, Blader IJ. <span class=“named-content genus-species” id=“named-content-1”>Toxoplasma gondii</span> Infections Alter GABAergic Synapses and Signaling in the Central Nervous System. mBio. 2015;6(6):e01428–15. doi: 10.1128/mBio.01428-15.

54. Boillat M, Hammoudi PM, Dogga SK, Pagès S, Goubran M, Rodriguez I, et al. Neuroinflammation-Associated Aspecific Manipulation of Mouse Predator Fear by Toxoplasma gondii. Cell Rep. 2020;30(2):320-34.e6. Epub 2020/01/16. doi: 10.1016/j.celrep.2019.12.019. PubMed PMID: 31940479; PubMed Central PMCID: PMCPMC6963786.

55. Poller WC, Downey J, Mooslechner AA, Khan N, Li L, Chan CT, et al. Brain motor and fear circuits regulate leukocytes during acute stress. Nature. 2022. doi: 10.1038/s41586-022-04890-z.

56. Goerner AL, Vizcarra EA, Hong DD, Bergersen KV, Alvarez CA, Talavera MA, et al. An ex vivo model of <em>Toxoplasma</em> recrudescence. bioRxiv. 2020:2020.05.18.101931. doi: 10.1101/2020.05.18.101931.

